# The Arabidopsis TIRome informs the design of artificial TIR (Toll/interleukin-1 receptor) domain proteins

**DOI:** 10.1101/2025.03.15.643477

**Authors:** Adam M. Bayless, Lijiang Song, Mitchell Sorbello, Sam C. Ogden, Tyler S. Todd, Alice Flint, Natsumi Maruta, Jedidiah Tulu, Mikhail Drenichev, Vardis Ntoukakis, Thomas Ve, Mehdi Mobli, Li Wan, Qingli Liu, Jeffery L. Dangl, Bostjan Kobe, Murray Grant, Marc T. Nishimura

## Abstract

The TIR (Toll/interleukin-1 receptor) domain is an ancient protein module that functions in immune and cell death responses across the Tree of Life. TIR domains encoded by plants and prokaryotes function as enzymes to produce diverse small molecule immune signals. Plant genomes can encode hundreds of TIR-domain containing proteins – many of which confer important agricultural disease resistance as TIR-NLR (nucleotide-binding, leucine-rich repeat) immune receptors. Despite their importance, how natural variation influences TIR enzymatic output and immunity-associated cell death is largely unexplored. We assayed a complete collection of the TIR domains of *Arabidopsis thaliana* Col-0 (the “AtTIRome”) to explore variation in TIR metabolite production and cell death signaling. Roughly half of the AtTIRome triggered cell death in transient assays. Artificial TIR proteins designed based on consensus sequences of the AtTIRome’s cell death phenotypic classes revealed polymorphisms controlling variation in TIR cell death elicitation and metabolite production. Structure-function analyses of artificial TIRs revealed that natural variation in the “BB-loop”, a flexible region overlying the catalytic pocket, determines differences in function across Arabidopsis TIR-containing proteins. We further demonstrate that artificial TIRs are functional on an NLR chassis and that BB-loop variation can tune the activity of a natural TIR-NLR protein. These findings shed light on the diversity of TIR outputs and reveal methods to design and engineer TIR-based immune receptors.

**Significance Statement:** TIR (Toll/interleukin-1 receptors) domain proteins perform immune signaling across the Tree of Life. TIR domains are enzymes that can process NAD^+^ to generate diverse small molecule signals. In order to better understand TIR-based signaling, we leveraged Arabidopsis natural variation across ∼150 TIR proteins to inform the design of artificial TIRs, define features that control output, and tune the output of a natural TIR immune receptor. The engineering of plant immune pathways will enable the optimization of disease resistance to safeguard yields.

## Introduction

Humanity is projected to require 40% more food by 2050, yet around 15-20% of crop yields are lost each year to pests and disease (1, 2). Enhancing plant immune system function can help limit losses, and recent advances in genome editing and protein engineering offer to complement traditional plant breeding efforts to safeguard food production (3–5). Many plant disease resistance traits are conferred by TIR-NLR (Toll/interleukin-1 receptor, nucleotide-binding site leucine-rich repeat) intracellular receptor proteins (6, 7). TIR domains are evolutionarily ancient and often function in prokaryotic and eukaryotic immune systems (7–11). In plants and bacteria, TIR domains act as enzymes to produce diverse small molecules that can activate immunity and cell death. A better mechanistic understanding of TIR functions should enable the rational engineering of plant immune systems and allow for predictable tuning of signal outputs to boost resistance or reduce the agronomic costs of auto-activity.

The plant immune system can sense and respond to extracellular microbial patterns (pattern-triggered immunity), as well as to intracellular virulence factors (effector proteins) injected into plant cells by pathogens (effector-triggered immunity, or ETI) (6, 12). Often, ETI activation results in a localized host cell death known as the hypersensitive response (HR) that limits pathogen spread. While NLR-based ETI disease resistance is a valuable agronomic trait, in some cases the costs of mis-regulated immune activation can be high (13, 14). Mis-regulation of innate immune receptors is particularly likely to occur when immune receptors are moved between genomes, or when engineered to achieve new specificities (15). Thus, rational engineering of NLR innate immunity could benefit from generalizable solutions to re-regulate immune outputs.

Plant NLRs function in the cytoplasm as multi-domain switches to activate innate immunity (16). The canonical NLR has an N-terminal signaling domain, a central NBS (nucleotide binding site) domain, and a C-terminal LRR (leucine-rich repeat) domain. NLRs are typically activated when the C-terminal LRR domain recognizes pathogen effectors and/or their activities (12). Once activated, conformational changes in the central NBS domain allow the NLR to oligomerize into a ‘resistosome’, which promotes immune signaling by induced proximity of the N-terminal TIR or CC (coiled coil) domains (17–20). Similar to mammalian inflammasomes, plant resistosomes are wheel-like structures composed of four to six protomer NLR subunits (21–23). While CC domains can directly transduce signals by oligomerizing into ion channels, TIRs signal by oligomerizing to engage their intrinsic enzymatic activities and generate small molecule signals (24–31).

Enzymatic plant TIR domains convert nucleotide-containing substrates like NAD^+^ (nicotinamide adenine dinucleotide), NTPs (nucleoside triphosphates), and DNA or RNA either directly or indirectly into a variety of metabolites, including pRib-AMP/ADP (phosphoribose-adenosine monophosphate/diphosphate), ADPr-ATP (ADP-ribosylated ATP), ADPr-ADPR (di-ADPR), 2’cADPR (cyclic ADP-ribose), 2’,3’-cNMP, (cyclic nucleotide monophosphate), and RFA (ribofuranosyladenosine) (17, 20, 25, 30–34). The enzymology underlying the production of these diverse molecules by TIR domains is poorly understood, but requires a conserved putative catalytic glutamate and oligomerization-dependent coordination of a nearby “BB-loop” motif. (7, 35). The best characterized TIR-generated signals, pRib-AMP/ADP and ADPr-ATP/di-ADPR, are relayed by EDS1 (Enhanced disease susceptibility 1) family proteins into ETI outputs (30, 31). After binding TIR metabolites, EDS1 family protein complexes activate the downstream helper NLR (hNLR) proteins ADR1 (Activated disease resistance 1) or NRG1 (N requirement gene 1), which oligomerize into ion channels to promote transcriptional defenses and localized host cell death (26, 27, 32, 33, 36–38). pRib-AMP/ADP and ADPr-ATP/di-ADPR are apparently low abundance and/or unstable molecules whose detection is presently limited to cryo-EM or LCMS analysis after capture and protection by an in vitro-purified EDS1 complex (30, 31). Other TIR-produced metabolites (2’cADPR, 3’cADPR, 2’,3’cNMP, and RFA) are more easily detectable, but their specific roles as plant signaling molecules remain to be firmly established (7, 25, 32, 39, 40). RFA and 2’cADPR are structurally similar to the EDS1 immune signal pRib-AMP, and are readily detected *in planta* (34, 39, 41). 2’cADPR can be hydrolyzed to pRib-AMP by plant extracts and is a plausible precursor or storage form of this EDS1-activating signal (Yu, et al., 2024). Certain plant pathogens induce RFA accumulation (41, 42), but only recently was RFA characterized as a biomarker of enzymatic TIR activities (34). In the prokaryotic TIR-based Thoeris innate immune system, 3’cADPR functions as a signal to activate antiphage defense, and both 2’ and 3’cADPR are produced by bacterial plant pathogen virulence proteins inside host cells (29, 39, 43–45). How these diverse TIR-produced small molecules regulate plant-pathogen interactions remains a major unanswered question.

Plant TIRs require a conserved catalytic glutamate for catalysis, as well as conserved oligomerization interfaces (the “AE” and “BE” interfaces), yet TIR domains within a genome can share less than 40% identity and occur in various domain architectures aside from canonical TIR-NLRs (7, 25, 46–48). Because dicot plants frequently encode hundreds of different TIR domain-containing proteins, an understanding of how TIR diversity influences their enzymatic profile (product types, abundance), and in turn, immune outputs, will be critical to successfully engineer crop TIR signaling and design customized TIR immune receptors (4, 49–52).

Plant pathogens continually evolve virulence effectors that evade or overcome recognition by host LRR domains of NLR receptors (53–56). Similarly, LRR domains of NLRs have been engineered to restore or expand effector detection (4, 5, 57–60). Recently, engineering of the central NBS domain has been shown as a viable strategy to resurrect defeated NLRs (5). However, the engineering of NLR signaling domains (TIR or CC) - to heighten or dampen immune outputs - is largely unexplored.

Relatively few TIR proteins have been studied, and the vast majority of plant TIR diversity remains unexamined (61). Here, we survey TIR natural variation within the *Arabidopsis thaliana* Col-0 genome and characterize the metabolite and *EDS1*-dependent cell death output of 148 TIR domains (the “AtTIRome”). We then leverage the AtTIRome to design artificial TIR proteins with distinct profiles of metabolite production and cell death-triggering function. Finally, we transfer a motif revealed by artificial TIR functions back into the full-length TIR-NLR, RPP1 (*Resistance to Peronospora parasitica 1*), to demonstrate the tuning of effector-activated TIR-NLRs outputs. These studies suggest that tuning of TIR domains at the BB-loop may be a generalizable strategy to re-regulate immune receptors.

## Results

### Identification of active TIR domains in the TIRome of *Arabidopsis thaliana* (Col-0)

To understand how natural variation influences TIR signaling, we screened each *Arabidopsis thaliana* (Columbia; Col-0) encoded TIR (the AtTIRome, 148 TIRs) for cell death induction and metabolite production in *Nicotiana benthamiana* (Fig. 1A and B). The AtTIRome includes TIR proteins with diverse architectures, including TIR-NBS-LRR, TIR-NBS, TIR-only, TIR-TIR, and xTNx/TNP (TIR-NBS/ARC-tetratricopeptide-like repeat). Plant TIR-NLRs are typically activated by effector-triggered oligomerization, but the specific effector triggers for most *Arabidopsis* TIRs are not known. Given these constraints, we activated the Arabidopsis TIR domains via constitutive oligomerization, enabling us to focus on the intrinsic properties of the TIR domains themselves. Plant TIR domain oligomerization and activation can be artificially driven by fusion to the SAM (sterile alpha motif) domain of the human TIR protein, SARM1 (sterile alpha and TIR motif containing 1) (24, 25, 35, 39, 62). SAM-TIR proteins retain a functional requirement for the native TIR self-association interfaces, the catalytic glutamate (E) residue, and still signal through EDS1, indicating that SAM-TIRs at least partially mimic TIR-NLR activation (24, 25). We fused HA-tagged SAM domains on each Arabidopsis TIR to assess their activity and protein accumulation (via immunoblot) in *Nicotiana*. In most cases, the initiating methionine defined the N-terminus of the TIR ORF (open reading frame) clones. TIR domains of multidomain proteins (*e.g.* TIR-NLR and TIR-NBS) were typically C-terminally truncated immediately upstream of the easily identifiable conserved Walker A motif (GxxxxGK[S/T]) in the NBS domain. This C-terminal truncation site was chosen based on the observation that the TIR-NBS “TIR+80” linker is required for full auto-activity of the RPS4 TIR (25, 63).

**Figure 1.**
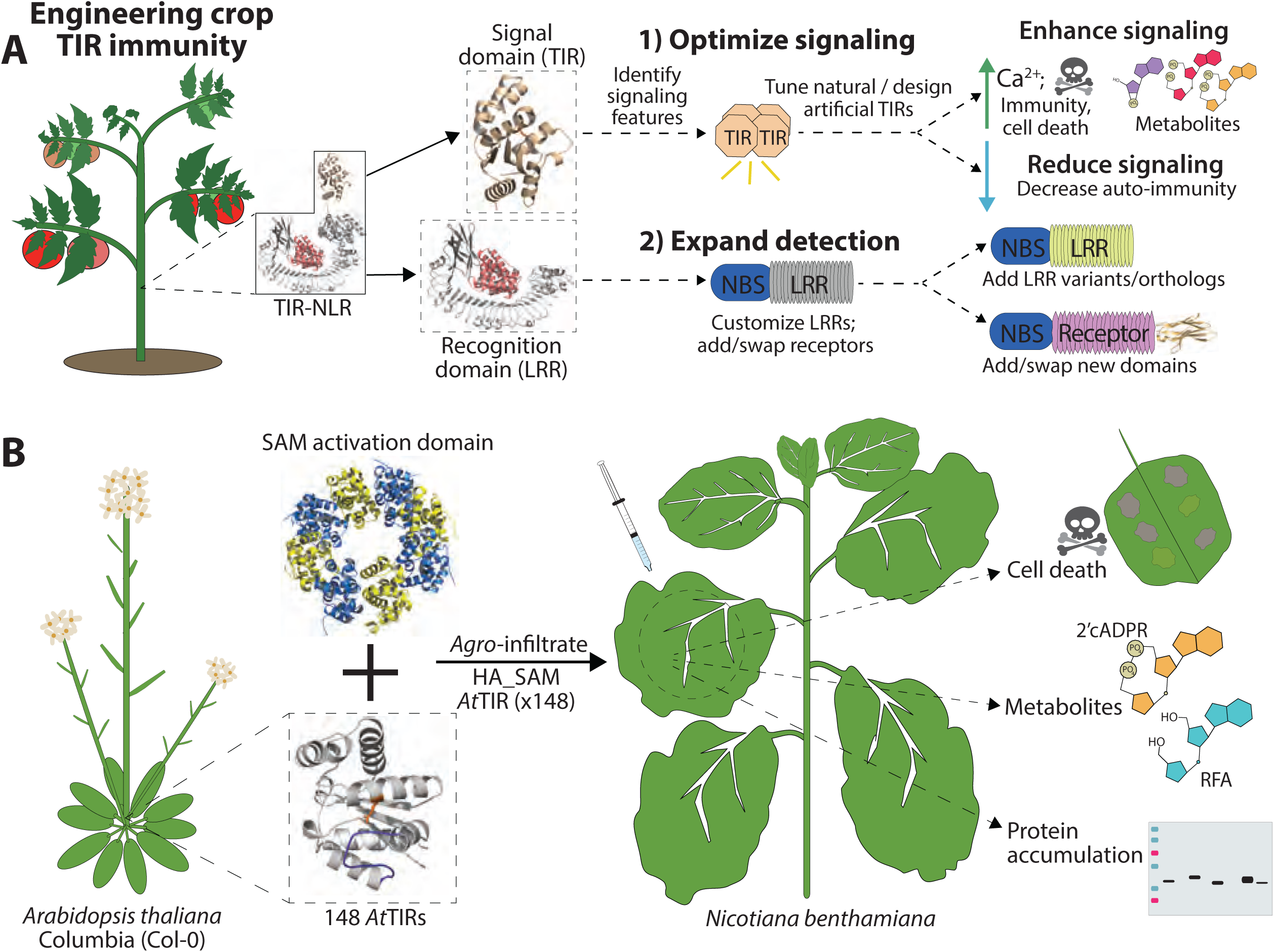
Strategies to engineer plant TIR-NLR immune receptors and screen natural diversity. (*A*) Engineering plant TIR-NLR detection (LRR) or signal domains (TIR) to alter recognition or optimize signaling. (*B*) Approach to phenotype each *Arabidopsis thaliana* (Col-0) encoded TIR domain (the AtTIRome) using *Nicotiana benthamiana* (*Nb*). Each AtTIR was fused to a HA_SAM domain from SARM1 (sterile alpha and TIR-motif containing 1; PDB ID 6ZGO), which promotes TIR activation, and screened for cell death induction, metabolite production, and protein accumulation after 35S transient expression via *Agrobacterium* infiltration.

The core immunity mediator EDS1 relays TIR metabolite signals into outputs such as localized cell death (7, 17, 25, 52). Accordingly, we scored each AtTIR for the ability to trigger EDS1-dependent cell death. *Agrobacterium tumefaciens* delivering each SAM*-*TIR construct was infiltrated into leaves of wild-type (WT) *Nicotiana benthamiana* (*Nb*) and scored for visible cell death (Fig. 1B, Fig. 2, SI 1). Approximately 47% (70/148) of the AtTIRome consistently activated EDS1-dependent cell death, as defined by necrosis or chlorosis. As expected, none of the 15 AtTIRs that lacked a catalytic glutamate (E) residue triggered auto-active cell death. Roughly 43% of AtTIRs had a catalytic E but did not trigger auto-active cell death (Fig. 2, SI 1). We also examined the accumulation of each SAM-TIR protein using *Nb eds1* plants, as EDS1-dependent cell death could increase TIR turnover and confound interpretation (See SI 2 for immunoblots). A strict correlation between protein abundance and cell death induction was not apparent – importantly, some strongly accumulating TIRs did not signal cell death and *vice versa* (Fig. 2, SI 1, SI 2). Many AtTIRs from atypical architectures like TIR-PP2 (phloem protein domain) or TIR-TIR (‘dumbbell’ TIRs) also signaled cell death, revealing that SAM fusions can activate diverse TIR domains from non-TNL architectures.

**Figure 2.**
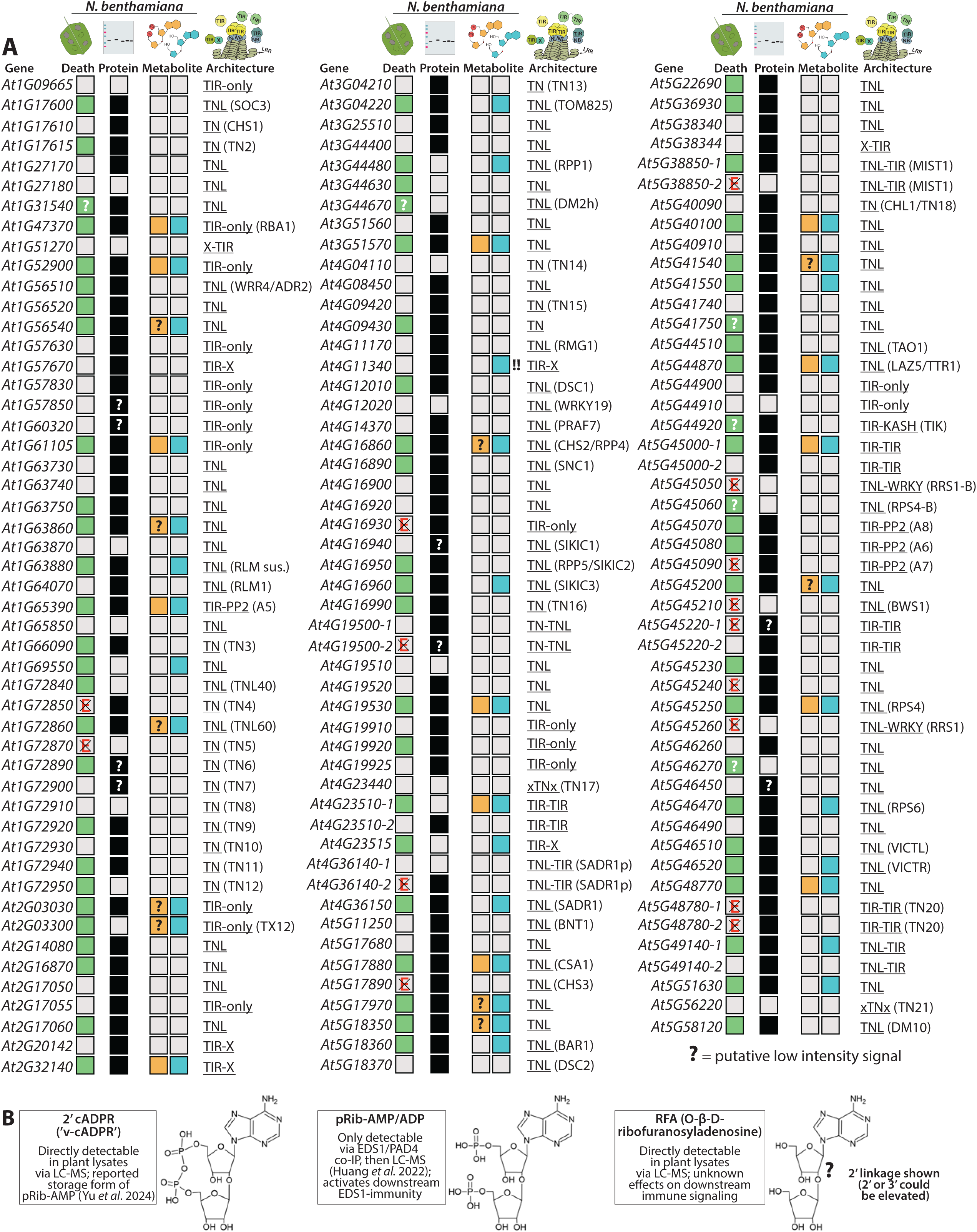
The *Arabidopsis thaliana* Col-0 TIRome includes diverse cell death and metabolite production phenotypes. (*A*) Cell death, protein accumulation, and metabolite production (2’cADPR, RFA (ribofuranosyladenosine)) by the AtTIRome (148 TIRs). A positive metabolite score is indicated by a filled box (orange for 2’cADPR, blue for RFA) and represents, minimally, a 3-fold elevation above that detected in control tissue. A crossed “E” indicates that the TIR lacks the conserved catalytic glutamic acid. The native genomic architecture of each AtTIR, and the gene name (if described) is provided. All AtTIRs were expressed as fusions to N-terminal HA_SAM domains and cell death was assayed in *Nb* WT plants. *Nb eds1* plants were used to assess metabolite production and protein accumulation. Cell death assays were performed at least three times. (*B*) Drawings of the TIR metabolites 2’cADPR (previously ‘variant cADPR’ or ‘v-cADPR’), pRib-AMP, and RFA. 2’cADPR and RFA are structurally similar to pRib-AMP. 2’cADPR may be a precursor or storage form of pRib-AMP, but is not known to directly signal via EDS1 (30, 31, 33).

Most tested plant TIRs are dependent on EDS1 to trigger resistance and auto-active cell death (64). Curiously, in our assays, several TIR domains (from *At2G32140*, *At3G51570*, *At4G16860*, *At5G45250* (*RPS4*) and *At5G48770*) caused chlorosis independently of *EDS1*, suggesting they could participate in non-canonical TIR pathways, or potentially cause NAD^+^ depletion similar to SARM1 in animals (Fig. 2, SI 1) (39, 62). A monocot TIR protein of the xTNx (or TNP) class was recently proposed to function independently of EDS1; however, in that study no dicot xTNx proteins were reported to cause cell death (64, 65). Consistent with this, we did not observe cell death driven by the TIR domains of two *Arabidopsis* xTNx encoding genes (*At5G56220* and *At4G23440*) (SI 1).

Nearly half of the 148 AtTIRs signaled EDS1-dependent cell death, despite encompassing a wide range of sequence diversity (See SI 3 for a phylogenetic tree). To understand differences between TIRs that did, and did not, activate cell death, we split the AtTIRs into two classes based on their cell death signaling phenotype: ‘Death’ or ‘Non-death’ (hereafter ‘Non’). We next assessed TIR domain metrics for the classes such as TIR-TIR interface conservation, BB-loop length/electrostatic charge, TIR domain length, and lengths outside the core TIR domain (SI 4).

Among the ‘Death’ group, the BB-loop was typically 17-19 residues, and previously described conserved residues were often present at the AE- and BE-interfaces (e.g. SH/SF and G residues, respectively), while the ‘Non’ group was more variable for either metric (25, 66). However, we did not detect substantial differences in predicted core TIR domain length, BB-loop electrostatic charges, or lengths outside the core TIR domain (SI 4). The above TIR regions were delineated according to multiple sequence alignment (MSA) and mapping onto the RPP1 (*Resistance to Peronospora parasitica 1*) TIR-NLR structure, a subset are shown as examples in (SI 4) (17).

As noted above, plant TIRs produce diverse metabolites including 2’cADPR, 2’,3’-cNMP, RFA, pRib-AMP/ADP, and ADPr-ATP/di-ADPR (7, 30–32). pRib-AMP/ADP and ADPr-ATP/di-ADPR activate EDS1-signaling; however, no methodologies to detect these molecules directly within plant extracts have been reported (30, 31). As 2’cADPR can be metabolized to pRib-AMP, it is a plausible proxy for this EDS1-activating signal (33). Therefore, we assessed AtTIR enzymatic activity via LC-MS detection of 2’cADPR and ribofuranosyladenosine (RFA) within *Nb eds1* leaf extracts (Fig. 2; chromatographs shown in SI 5). The analytic method used is unable to discriminate between 2’RFA and 3’RFA, so our usage of “RFA” is agnostic as to the ribose-ribose linkage (34). Three different metabolite profiles were observed among the 70 AtTIRs that consistently signaled EDS1-cell death: RFA-only, RFA and 2’cADPR, or none detected. 53% (37/70) elevated RFA, however, only 20% (14/70) generated 2’cADPR above background levels. Notably, any AtTIR that produced 2’cADPR also generated RFA. The other 47% (33/70) of death signaling TIRs did not elevate RFA (or 2’cADPR) above background. This suggests that these 33 TIRs possess a weaker catalytic activity, which nonetheless, is still sufficient to stimulate EDS1 (potentially via catalysis skewed towards ADPr-ATP/di-ADPR). Although all LC-MS profiling included transitions for pRib-AMP/ADP and ADPr-ATP/di-ADPR, these were not detected, indicating that they were absent or below the limit of detection. 3’cADPR was not detected in any of the transient assays. The metabolite 2’,3’-cAMP was consistently not elevated among the initial batches of AtTIRome samples (which included both cell death positive and negative classes), and so was not measured in the remaining samples (SI 6). 2’,3’-cNMP production by TIR proteins has only been reported in the context of TIR-only proteins or isolated TIR domains; therefore, it is unclear if our addition of an octameric SAM domain might limit the formation of the indeterminate-length TIR filament structures associated with 2’,3’-cNMP generation (32).

### The AtTIRome can inform the design of functional artificial TIR proteins

Given that the AtTIRs could be divided into phenotypic classes, we next attempted to leverage TIR diversity to better understand structural motifs associated with TIR function across the AtTIRome. Particularly, we asked if our AtTIRome phenotypic classes could inform the design of artificial TIR proteins with predictable phenotypes (Fig. 3A). We determined the consensus sequence of the AtTIRome groups, ‘Death’ and ‘Non’, as well as the consensus of all 146 AtTIRs (hereafter ‘All’). Any sequences outside of the predicted core TIR domain were trimmed, and the two xTNx TIRs were excluded, as they are proposed to be mechanistically distinct (64). The resulting artificial consensus TIRs for ‘Death’, ‘All’, and ‘Non’ were 160-162 residues in length each. AlphaFold3 models indicate that the artificial consensus TIRs had catalytic E residues, canonical TIR-TIR interfaces, and had similar residues within the putative catalytic pocket proposed by Manik *et al* (29) (SI 7). Given this conservation of TIR features, the ‘Non’ consensus was outwardly similar to enzymatically active TIRs and lacked predictable loss of function polymorphisms. Figure 3B provides AlphaFold predictions for each artificial consensus TIR, as compared to the RPP1 TIR domain structure (confidence scores in SI 7) (17). The majority of variation among the artificial consensus TIRs was found at the BB-loop or the αC and αD helices, which are located away from TIR-TIR interaction interfaces (see SI 8 for sequence alignment).

**Figure 3.**
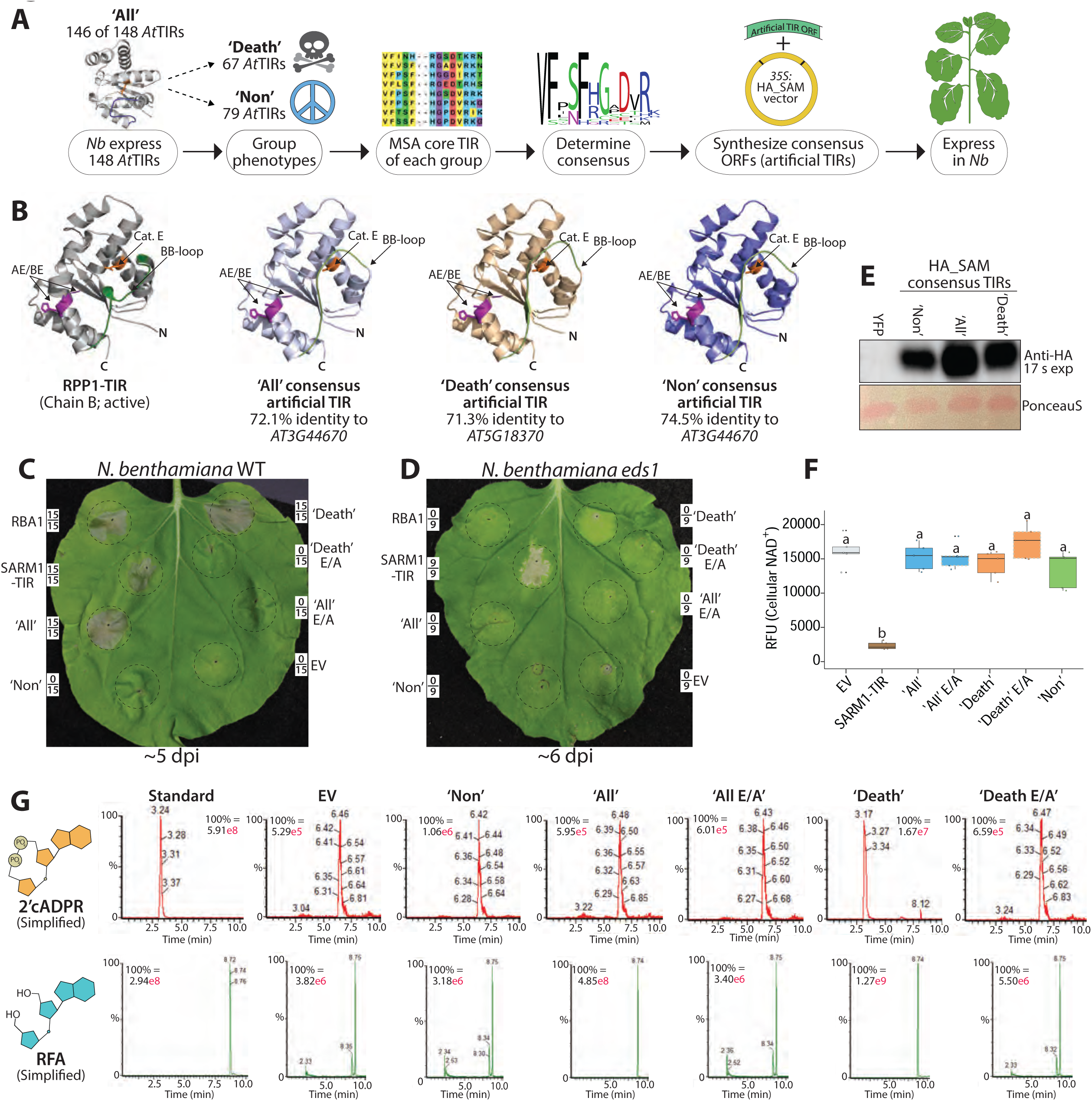
Artificial TIRs designed from AtTIR consensus groups are enzymatically active and signal cell death. (*A*) Diagram of artificial TIR design using AtTIR consensus group sequences. (*B*) (Left) Structure of an active state (chain B) TIR domain from the TIR-NLR RPP1 (PDB 7CRC) (17); BB-loops are colored green, catalytic glutamates are shown in orange, and AE/BE interfaces are shown in purple. (Center-Right) AlphaFold structure predictions of the three artificial consensus TIRs, ‘All’, ‘Death’ or ‘Non’. N: N-terminus, C: C-terminus. (*C, D*) *Nb* WT or *eds1* leaves expressing artificial TIRs with HA_SAM fusions. Control RBA1, SAM_SARM1-TIR (residues 478-724) or empty vector (EV; 35S:YFP) were similarly expressed. Leaves are shown ∼4-5 days post-infiltration, and framed numbers denote replicates per set. (*E*) An anti-HA immunoblot showing artificial TIR accumulation in *Nb eds1* leaves at 40 hpi. (*F*) Fluorescence-based NAD^+^-detection assay in *Nb eds1* leaves expressing artificial TIRs or SARM1-TIR; assay performed at 40 hpi. RFU: relative fluorescence units. TIR domain catalytic glutamate mutants have alanine substitutions and are noted as (E/A). (*G*) LC-MS traces of 2’cADPR or RFA from artificial TIRs in *Nb eds1* leaves at 40 hpi. Similar experiments were performed at least three times. Statistical analyses: one-way ANOVA and Turkey HSD with CLD (compact letter display) of significance classes. Over-lapping letters are ns (non-significant; *p*>0.05) and a separate letter class indicates *p*<0.05 or better.

Next, we synthesized ORFs encoding the ‘Death’, ‘Non’ or ‘All’ artificial TIRs, and cloned them into SAM constructs, as used to screen the AtTIRome. In agreement with their class phenotypes, both ‘Death’ and ‘All’ triggered EDS1-dependent and catalytic E-dependent cell death in *Nb*, while ‘Non’ did not (Fig. 3C, D). The artificial consensus TIR proteins all accumulated protein by immunoblot (Fig. 3E). Curiously, ‘Death’ caused a minor chlorosis in *Nb eds1* plants, which required the catalytic glutamate (SI 9). Because cellular NAD^+^-depleting TIRs like SARM1 or AbTir cause EDS1-independent cytotoxicity, we examined if ‘Death’ might also diminish NAD^+^ levels in *Nb eds1* plants (Fig. 3F) (39). However, unlike SARM1-TIR, ‘Death’ did not deplete NAD^+^, indicating that the apparent chlorosis is NAD^+^ depletion-independent, or may be triggered by NAD^+^ reduction below the limits of detection (Fig. 3D). Despite being divergent at only 15 residues, the artificial consensus TIRs recapitulated the cell death phenotype of their class constituents and provide testable hypotheses on how natural variation might influence TIR signaling.

Because plant TIRs activate EDS1 via production of small molecule signals, the ‘All’ and ‘Death’ artificial TIRs were likely active enzymes. Accordingly, we used LC-MS to profile their measurable metabolic outputs (2’cADPR, RFA) (Fig. 3G). Similar to many AtTIRs in Fig. 2, ‘All’ elevated RFA but not 2’cADPR, while ‘Death’ elevated both 2’cADPR and RFA, as well as an unknown peak (*m/z* 542) at retention time ∼8.10 minutes (Fig. 3G). Metabolite elevation by either ‘All’ or ‘Death’ was dependent upon the catalytic glutamate (Fig. 3G). We also assayed TIR-only versions of ‘All’ and ‘Death’ that lacked SAM domains, and both generated metabolites and signaled EDS1-dependent cell death, indicating that their function does not necessarily require SAM-enforced oligomerization (SI 9). ‘Non’ did not activate EDS1-dependent cell death, and as expected, it did not elevate metabolites relative to EV (empty vector) controls (Fig. 3G).

In addition to designing artificial TIRs by consensus, as a control, we tested whether artificial TIRs generated by randomly picking residues at each position (from a pool of 146 AtTIRs) might also be functional (SI 10). We synthesized 10 artificial ‘random TIRs’ ranging from ∼48%-58% identity to the closest natural occurring TIR (SI 10). Each random TIR had a catalytic E. When expressed as a SAM fusion in WT *Nb,* none caused apparent cell death or chlorosis (SI 10). Together, these findings suggest that a consensus-based approach can reliably design functional artificial TIR proteins.

### Variation in the BB-loop of artificial TIRs controls cell death signaling and enzymatic activity

Several TIR regions, such as the BB-loop, the TIR-TIR interfaces, and catalytic pocket, can impact self-association, enzymatic activity and signaling (24, 25, 48, 66, 67). The ‘All’ and ‘Non’ artificial TIRs are ∼90% identical, possess catalytic E residues, and have identical TIR-TIR interfaces (Fig. 3). Yet ‘All’ produces RFA and activates cell death, while ‘Non’ does not. An amino acid sequence alignment revealed variation at the BB-loop (Region I) and two alpha-helices (helices αC and αD; referred to as Regions II and III, respectively) (Fig. 4A, SI 8). SI 11 shows these variant regions in context of a TIR tetramer (RPP1 model). To understand which polymorphisms were sufficient to restore activity, we swapped each of the three ‘All’ regions, and combination of regions, into ‘Non’ (Fig. 4B, C, E, F). BB-loop (Region I) transfer from ‘All’ to ‘Non’ was sufficient to restore signaling and elevate RFA production (Fig. 4B, C, F); however, adding either helix αC or αD variation (Region II or III) did not restore function, nor enhance signaling or metabolite production, if combined with BB-loop transfer (SI 11). We then asked which variant residues within the ‘All’ BB-loop were necessary for functionality in the context of the Non TIR domain. Removing the Q or E residues from the BB-loop abolished cell death triggered by the ‘Non’ TIR domain containing the ‘All’ loop (Fig 4D). Similarly, adding back the ‘Non’-specific G residue also abolished cell death triggered in the same context (Fig 4D). The T/V substitution had no impact on cell death in the same context (Fig. 4D). Together, this indicates that multiple BB-loop polymorphisms are necessary to restore signaling to ‘Non’.

**Figure 4.**
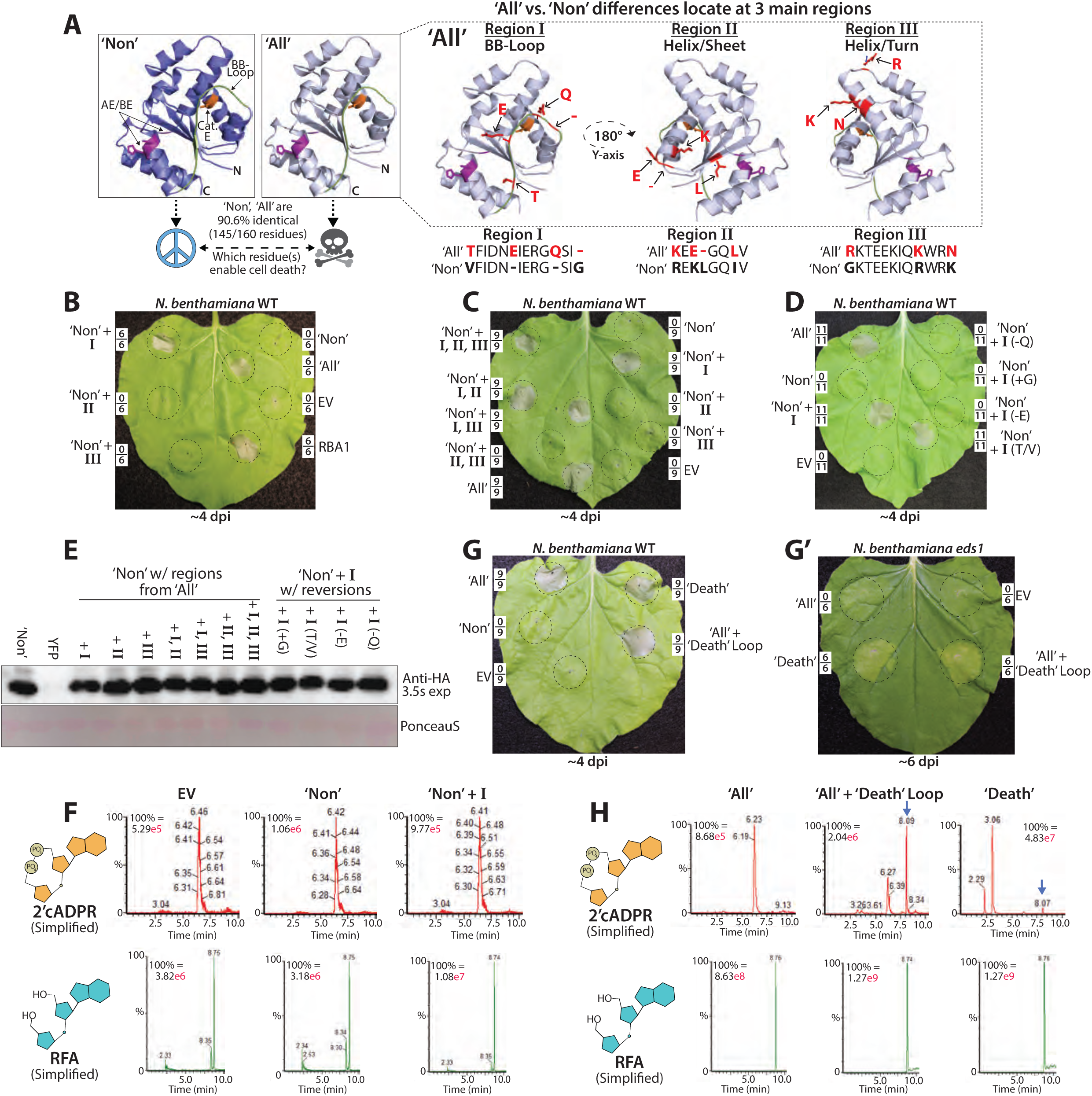
BB-loop variation determines cell death and metabolite production by the artificial consensus TIRs. (*A*) AlphaFold predictions of the artificial consensus TIRs, ‘All’ or ‘Non’, highlighting 11 of 15 variant residues. BB-loop colored green, catalytic glutamate shown orange, and AE/BE interfaces shown purple. Variant residues are colored red and indicated by arrows. N: N-terminus, C: C-terminus. (*B, C, D*) *Nb* WT leaves expressing ‘All’ or ‘Non’ artificial TIRs, as well as BB-loop or regional swaps. I, II, or III refers to a variant region in ‘All’ vs. ‘Non’. Leaves are shown ∼5 dpi and framed numbers denote replicates per set. (*E*) Anti-HA immunoblot of ‘All’, ‘Non’, and ‘Non’-variants from *Nb eds1* leaves at ∼40 hpi. (*F*, *H*) LC-MS traces of 2’cADPR or RFA from *Nb eds1* leaves expressing SAM tagged artificial TIRs and artificial TIR variants. Leaf samples harvested ∼40 hpi; EV (empty vector) refers to 35S:YFP. (*G*) *Nb* WT or *eds1* leaves expressing various artificial TIRs, including ‘All’ with a BB-loop swap from the ‘Death’. Leaves shown ∼5 dpi and framed numbers denote replicates per set.

We next examined if the BB-loop from the ‘Death’ artificial TIR modulated enzymatic outputs. As such, we replaced the loop of ‘All’ with that from ‘Death’ and detected a new peak at ∼8.10 min, as well as a gain of *EDS1*-independent chlorosis similar to that observed with the ‘Death’ TIR protein (Fig. 4H, G, G’; Fig. 3D). Collectively, these results suggest that the AtTIRs comprising the ‘Non’ group are enriched for BB-loops that are suboptimal for triggering cell death, and that BB-loop variation can influence TIR metabolic outputs.

### *In vitro* characterization of artificial TIRs

As noted above, methodologies to directly detect pRib-AMP/ADP or ADPr-ATP/di-ADPR *in planta* are lacking, so we also examined TIR outputs *in vitro*, using recombinant proteins. We first performed kinetic NADase assays and observed that ‘Non’ was inactive compared to the ‘All’ or ‘Death’ TIRs, and that, consistent with *in planta* assays, only ‘Death’ produced 2’cADPR (Fig 5A, SI 12A-C). Consistent with *in planta* assays in Figure 4, swapping the ‘All’ BB-loop into ‘Non’ promoted NAD^+^-hydrolysis, while placing the ‘Death’ BB-loop into ‘All’ enhanced the rate of NAD^+^ hydrolysis over 10-fold (Fig 5A). Each *in vitro* NADase timepoint was assessed using ^1^H NMR spectroscopy and compared with available standards (Nam, 2’cADPR, and NAD^+^) (see SI 12A for spectra). Interestingly, the 2’cADPR peak for ‘Death’ increased exponentially until 4 h, and then increased linearly despite no more NAD^+^ being present, suggesting this TIR might also bind and cyclize ADPR (SI 12B).

**Figure 5.**
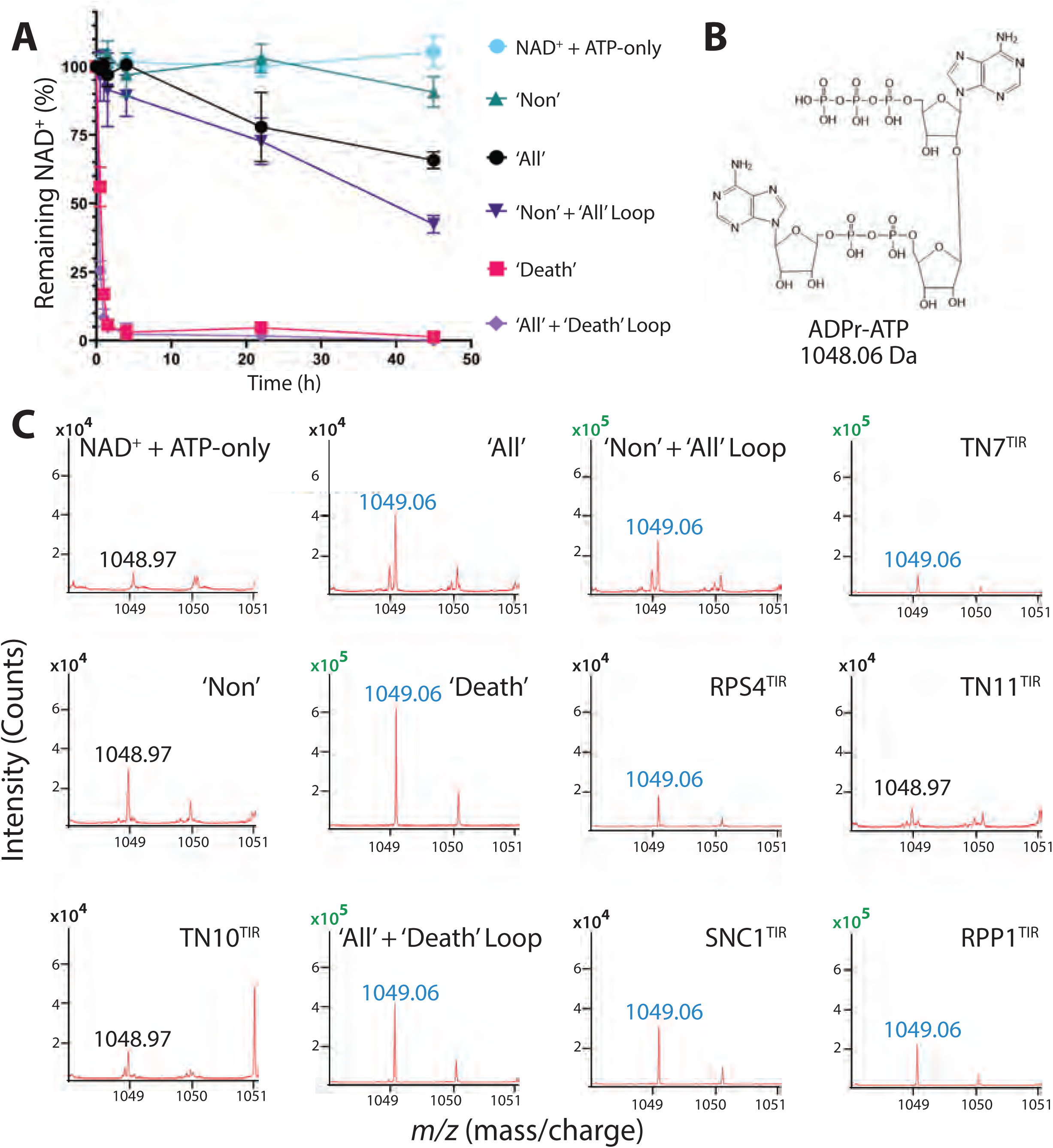
*In vitro* NADase assays of artificial and natural TIRs. (A) Kinetic NADase assay of recombinant artificial TIRs. The intensity of the signal corresponding to NAD^+^ at 0 h, measured via ^1^H NMR, was compared to the intensity remaining at the indicated time-points. The mean values of triplicates are shown with standard error bars. (*B*) Stick representation of the structure of the TIR metabolite ADPr-ATP; molecular mass 1048.06 Daltons. (*C*) MALDI-TOF MS detection of ADPr-ATP from *in vitro* TIR end-point reactions. Spectra for m/z regions 1048–1051 are shown. Blue (1049.06 *m*/*z*) corresponds to ADPr-ATP, while 1048.97 *m*/*z* (black) is present in the background control NAD^+^ + ATP and ‘Non’ spectra. Note: Y-axis shown either 10^4^ (black) or 10^5^ (green).

After establishing robust *in vitro* TIR assays, we examined ADPr-ATP production from endpoint reactions via MALDI-TOF (matrix assisted laser de/ionization – time of flight)-MS and ^1^H NMR (Fig 5BC, SI 12D-I). As controls, we included the RPP1 and RPS4 TIR domains - which are known to produce ADPr-ATP - as well as several TIR-only proteins, TN7, TN10, and TN11 (30, 31, 46). SI 12D shows NAD^+^ consumption for each examined TIR. Notably, ‘Non’ and TN10 did not accumulate Nam, consistent with a lack of NADase activity and being unable to trigger cell death *in planta* (SI 12D). All enzymatically active TIRs (except TN11) generated MS peaks matching predictions for ADPr-ATP (+H^+^) (1049.1 *m/z*) (Fig 5B, C). Curiously, all TIRs that produced ADPr-ATP peaks - as detected by MALDI-TOF-MS – also produced two unique peaks (U1, U2) detectable by ^1^H NMR (SI 12E, F). Although ‘Death’ was the only artificial TIR that could produce 2’cADPR, all surveyed Arabidopsis TIRs with enzymatic activity generated 2’cADPR (SI 12F). Together, the MALDI-TOF-MS and NMR spectra show that both natural and artificial TIRs can generate diverse metabolites *in vitro*, including EDS1-activating signals like ADPr-ATP.

Several active TIRs produced products with *m/z* 462.1 and 638.1 (SI 12G-K) in addition to 1049.1 (SI 12G-J). Interestingly, 638.1 *m/z* was observed within 1049.1 ms/ms spectra, while 462.1 and 638.1 share similar fragment ions to ADPr-ATP. Hence, 638.1 and 462.1 species could represent di-ADPR, pRib-ADP, or ADPr-ATP degradation. Standards of di-ADPR, ADPr-ATP, and pRib-ADP may help to explore these mystery species. Their biological relevance and varied abundance between tested TIRs remains to be understood.

### Artificial TIR variation can inform TIR-NLR engineering

Because plant NLRs couple effector detection to signal generation, we examined whether the artificial TIRs were functional if attached to a biologically-relevant NBS-LRR chassis (6, 12). We chose RPP1 as a model system, as it is well-understood, with a cryo-EM structure and genetic information on both recognized and unrecognized alleles of the pathogen effector ATR1 (ARABIDOPSIS THALIANA RECOGNIZED1) available (17, 68). We replaced the native TIR of RPP1-WsB (residues 85-248) with the ‘Non’, ‘Death’ or ‘All’ TIR, and co-expressed each in *Nb* with the cognate ATR1-Emoy effector, or a non-activating allele, ATR1-Emwa (68) (Fig. 6A, B, SI 13). Relative to RPP1 WT, the artificial TIR-NLRs showed reduced accumulation, although ‘Death’-NLR was more stable than ‘All’-NLR or ‘Non’-NLR. LC-MS analysis revealed that both ‘Death’-NLR and ‘All’-NLR’ elevated RFA, though ‘Death’-NLR was >10-fold higher. Curiously, ‘Death’-NLR did not elevate 2’cADPR, unlike expression as a SAM fusion or TIR-only protein (Fig. 6C, Fig. 3G, SI 9). Because ‘Death’-NLR generated >10-fold more RFA, we compared its cell death signaling to the ‘All’-NLR, where it signaled effector-dependent cell death in 100% of *Nb* leaves, in contrast to 12% for the ‘All’-NLR (Fig. 6B). These results indicate that artificial TIRs can be functional on an NBS-LRR chassis and have outputs largely consistent with those observed in the context of an orthologous SAM fusion.

**Figure 6.**
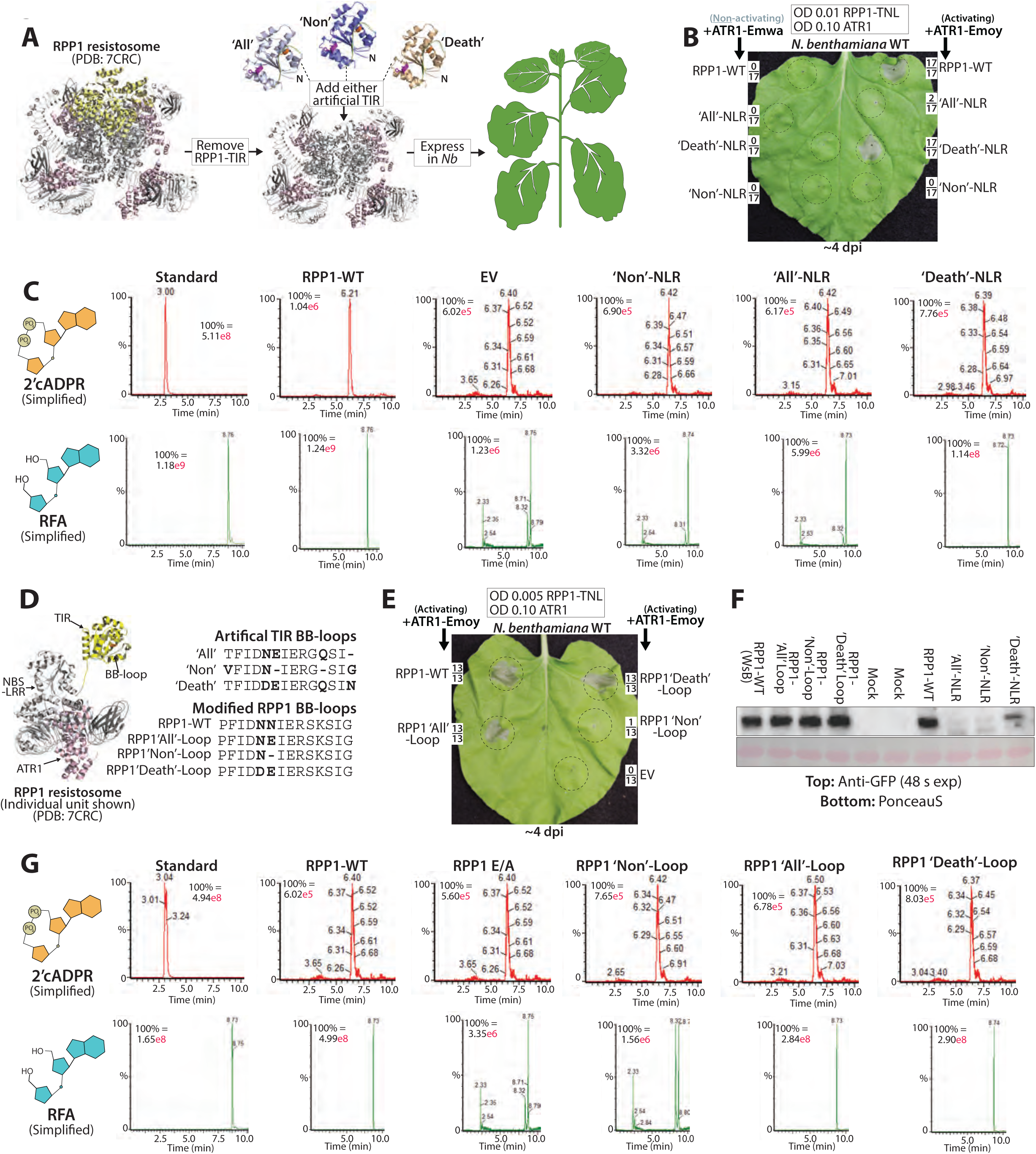
Incorporating BB-loop variation from artificial TIRs can modulate TIR-NLR signaling; artificial TIRs exhibit effector-mediated activation on NLR chassis. (*A*) Schematic diagram of the strategy to replace native RPP1 TIR domains with artificial TIRs. TIRs are shown in yellow, NBS-LRRs in grey, and ATR1 effectors in pink; PDB 7CRC (17). (*B*) *Nb* WT leaves expressing artificial TIR-NLRs at OD_600_ = 0.01. The cognate ATR1 (Emoy) allele triggers RPP1 WsB while the non-cognate Emwa allele does not. (*C*) LC-MS traces of 2’cADPR or RFA from *Nb eds1* leaves co-expressing artificial TIR-NLR fusions with the activating effector, ATR1 (Emoy). Tissue was harvested ∼40 hpi. (*D*) A monomer of the RPP1 TNL resistosome showing the TIR domain (yellow), NBS-LRR (grey), and activating ATR1 effector (pink). *Top*: Alignment showing artificial TIR BB-loop variation. *Bottom*: Modifying the RPP1 BB-loop to mimic artificial TIR variations. (*E) Nb* WT leaves expressing RPP1-WT (WsB allele) or RPP1-loop variants at OD_600_ 0.005; ATR1-Emoy co-expressed at OD_600_ 0.10. All RPP1 constructs contain C-terminal YFP tags, and EV refers to negative control 35S:YFP. Leaves were imaged ∼4 dpi and framed numbers denote replicates per set. (*F*) Anti-YFP immunoblot of RPP1-WT, RPP1 BB-loop variants, artificial TIR-NLRs, or ATR1 effectors (Emoy or Emwa). (*G*) LC-MS traces of 2’cADPR or RFA from *Nb eds1* leaves expressing WT RPP1, or RPP1 with BB-loop modifications, and the activating ATR1-Emoy effector.

We next asked whether variation within the artificial TIRs could inform TIR-NLR engineering. For example, might incorporating variation within the ‘Non’ BB-loop into a natural TIR-NLR dampen signaling? To do so, we replaced two BB-loop residues (N119, N120) of the TIR-NLR, RPP1, with the corresponding loop residues from ‘All’, ‘Non’, or ‘Death’ (Fig. 6D, E, G). When cognate ATR1-Emoy was co-expressed, we observed that RPP1_‘Non’-Loop_ triggered cell death in less than 10% of leaves, as compared to RPP1_‘Death’-Loop_, RPP1_‘All’-Loop_ or RPP1_WT_, (Fig. 6E). Similarly, RPP1_‘Non’-Loop_ did not elevate RFA, as compared to RPP1_WT_, RPP1_‘Death’-Loop_, or RPP1_‘All’-Loop_ (Fig. 6G). The RPP1 BB-loop variants accumulated similarly, as revealed by anti-YFP immunoblot, consistent with the ‘Non’-type loop polymorphisms affecting enzymatic activity (Fig. 6F). Thus, a BB-loop with reduced output discovered in the context of artificial TIR overexpression could inform tuning of native TIR-NLR activity in response to its cognate pathogen trigger.

## Discussion

TIR domains are conserved components of innate immune systems across the Tree of Life. In both prokaryotes and plants they function by generating diverse small molecule signals that activate downstream immune pathways (7, 47, 51, 52). To better understand the diversity of TIR domain function, we characterized each *Arabidopsis thaliana* (Col-0) TIR domain and found that differences in metabolite production (type, abundance) and cell death signaling are common. Further, we demonstrate that the Arabidopsis TIRome has sufficient information to enable the design of functional artificial TIRs, and that variation between the ‘Death’ and ‘Non’ classes can guide TIR engineering to tune natural TIR-NLR outputs in a predictable manner. Our results suggest that the BB-loop may be a useful generic target for controlling TIR-NLR activity and mitigating dysregulation resulting from NLR transfer between species or varieties.

Based on our current understanding of TIR pathways in plants, any cell death-signaling TIR should generate EDS1-activating metabolites like pRib-AMP/ADP or ADPr-ATP/di-ADPR (30, 31). Because current methodologies cannot directly detect these metabolites *in planta*, we measured 2’cADPR and RFA as biomarkers to assess TIR enzymatic activation (25, 34, 41, 42). Published literature contains conflicting results for 2’cADPR. Based on the structure, 2’cADPR was proposed as a plausible precursor of pRib-AMP (7, 29). However, early experiments expressing AbTir (a prokaryotic TIR that produces 2’cADPR) found that the resulting cell death was EDS1-independent, and thus a link to 2’cADPR was not supported (39, 40). More recently, Yu et al. found that AbTir can induce oligomerization of the *Arabidopsis* EDS1/ADR1 complex via pRib-AMP production and that 2’cADPR can indeed be hydrolyzed by plant extracts to pRib-AMP (33). While all active plant TIR domains produced 2’cADPR in our *in vitro* assays, we did not detect its cleavage into pRib-AMP. A full understanding of 2’cADPR’s function as a precursor or storage form of pRib-AMP remains to be established. While RFA is structurally similar to 2’cADPR and pRib-AMP, any signaling functions of RFA are unclear. RFA accumulates in *Arabidopsis* following *Pseudomonas syringae* DC3000 inoculation (41, 42). However, in these studies, RFA increases could reflect the enzymatic activity of host TIRs as well as that of the bacterial TIR effector HopAM1 (29, 39, 44). The TIR effector HopAM1 requires enzymatic activity for virulence and produces 3’cADPR, yet how 3’cADPR and/or manipulating host NAD^+^ mediates pathogenesis is not understood (29, 43, 44). 3’cADPR does not stimulate EDS1-mediated cell death, although 3’cADPR activates the Thoeris prokaryotic TIR immune system (29, 39, 40, 45). Our finding that RFA was detected more widely from active TIRs *in planta* than 2’cADPR reveals it as a useful biomarker, yet RFA was not elevated by ∼40% of the cell death signaling AtTIRs. It is plausible that some of the cell death-inducing TIRs that do not make detectable 2’cADPR or RFA are producing relatively more ADPr-ATP/di-ADPR to activate the SAG101/NRG1 branch of the EDS1 pathway. This will be important to understand as it might allow retuning of TIRs to signal preferentially for resistance via EDS1/PAD4/ADR or for cell death via the EDS1/SAG101/NRG1 branch (69). This possibly occurs for some natural TIRs such as TN11 (AT1G72940), which in our SAM-TIR system did cause cell death, and in our *in vitro assays,* exhibited NADase activity, produced 2’cADPR but not ADPr-ATP*I* Hence, the development of methodologies that can directly measure EDS1-activating signals in planta is essential to enable plant TIR research.

Almost half of the AtTIRome did not trigger cell death in *Nicotiana.* Does this represent the underlying biology (e.g. ADR vs NRG signaling as discussed above), or are there reasons that our screen might artifactually underrepresent AtTIRome signaling? There are several potential artifacts in our screen. For instance, we used auto-active SAM domain fusions to induce homo-oligomerization of the TIR domains, which might not effectively activate every TIR, particularly if hetero-oligomerization is required (70). Additionally, we noted substantial variation in SAM-TIR protein accumulation that might not reflect native TIR protein stability and/or accumulation. Given that certain non-accumulating SAM-TIRs could still trigger cell death, it is unclear to what extent we should expect protein accumulation to impact the phenotypes measured. For instance, while the TN7 (*At1g72900*) TIR produced ADPr-ATP *in vitro*, the SAM_TN7 TIR accumulated poorly *in planta* and did not trigger cell death. Regardless, non-enzymatic TIR-containing proteins could still fulfill valuable roles as sensors, similar to RRS1, and/or regulate the activities of tetrameric TIR complexes via hetero-oligomerization (17, 20, 70). Despite potentially underrepresenting TIR activity, our analysis of artificial TIRs in a transient expression system was able to pinpoint two residues in the BB-loop that predictably decreased activity of the natural TIR-NLR RPP1 after effector activation. This suggests that our approach, albeit artificial, was sufficiently robust to provide useful information about natural TIR-NLR receptor function.

Our findings add to a growing list of reports illustrating BB-loop influences on enzymatic TIR signaling (24, 25, 39, 67). The artificial consensus TIRs ‘All’ and ‘Non’ have identical TIR-TIR interface and catalytic residues, yet BB-loop differences prevent signaling by ‘Non’. The *in vitro* enzymatic activity of the RUN1 TIR domain can be enhanced by adding positive arginine residues into the BB-loop (24), yet the ‘Death’ TIR is enriched for negative BB-loop charges and transferring the ‘Death’ TIR BB-loop into ‘All’ increased enzymatic activity. BB-loop alterations can have different impacts on plant and prokaryotic immune TIRs. For instance, shortening and replacing several BB-loop residues within the prokaryotic ThsB TIR enables auto-active signaling, whereas similar replacements within auto-active plant TIRs abolish activity (39). A recent study by Song *et al* indicates that the BB-loop and TIR-TIR association interfaces can contribute to the formation of TIR condensates (Song et al., 2024).

The artificial consensus TIRs ‘All’, ‘Non’, and ‘Death’ had distinct metabolite and signaling profiles. ‘All’ only elevated RFA, while ‘Death’ produced RFA, 2’cADPR and an unknown peak at ∼8.1 min. Curiously, when ‘Death’ was fused to the RPP1-NLR, it no longer produced 2’cADPR or the 8.1 min peak, unlike TIR-only or SAM fusion contexts. Such context-dependency requires further investigation. Isolated TIR domains are reported to generate 2’,3’-cNMP, but how domain architecture might influence metabolite production is not understood (32). The activation of artificial TIR-NLRs by cognate effectors suggests that other ‘customized’ TIRs could likely perform regulated signaling upon an NLR chassis. Given that the RPS4 TIR can signal as a NLRC4 chimera suggests that artificial TIRs could likely function on atypical recognition domains, including customized LRR-nanobody detection domains (‘pikobodies’) (40, 57).

While we designed artificial TIRs using a consensus approach on a mesoscale set of ∼150 TIR domains, artificial proteins of various families can also be generated using large language models (LLMs) trained on millions of protein sequences (3, 71, 72). For instance, Madani *et al.* produced functional artificial lysozyme proteins with identities as low as 31% to WT HEWL (hen egg white lysozyme), although artificial lysozyme proteins with identities closer to 70% of WT displayed higher activity (3). Resources like the OpenPlantNLR initiative, which houses ∼60,000 NLR sequences - including thousands of TIRs from various species - could help design new artificial TIRs (https://zenodo.org/communities/openplantnlr/). Similarly, a large set of species-specific TIRs as in the *Arabidopsis* pangenome could provide considerable design power (61). The compact size of TIR domains (∼160 residues) allows economic large-scale screening of natural TIR domains and artificial TIR designs. While thousands or millions of sequences are available for many protein families, phenotyping proteins remains a bottleneck. Our consensus-based approach indicates that smaller scale studies, employing low/medium-throughput in vivo phenotyping, can be informative.

Although this study demonstrates the production of artificial immune signaling domains, key questions remain about plant TIR engineering. For instance, which TIR variations dictate substrate usage, or the production of specific metabolites? How do different TIR signals translate to actual crop success, and is yield benefited by TIRs with broad (2’-cADPR, RFA, 2’,3’-cNMP, pRib-AMP/ADP, ADPr-ATP/di-ADPR) or more narrow metabolite outputs? And can similar engineering principles be applied to TIRs that reportedly function outside of EDS1-signaling pathways (34, 64)?

Enzymatic TIRs that make unusual signaling molecules continue to be discovered (73–75). Prokaryotes encode diverse assortments of enzymatic TIRs, and a mechanistic study of these should offer further insights into how particular TIR regions and their overall context in a protein complex impart product and/or substrate specificities (8, 9, 11, 29, 43, 44, 76). Artificial protein-based approaches may also facilitate the design of immune proteins belonging to other families, such as those containing CC domains. As our mechanistic understanding of plant immune receptors increases, protein engineering offers to complement traditional crop improvement techniques with rationally engineered disease resistance.

## Methods

### Gene Cloning & DNA Vector Construction

*Arabidopsis thaliana* accession Col-0 TIR ORFs (AtTIRs) were cloned from cDNA (or synthesized) beginning at the native start codon (ATG) to the start of the P-loop. All AtTIR ORFs contained BP-compatible recombination ends and were placed into pUC57-Kan or pDONR207. Artificial TIRs were codon-optimized for expression in *N. benthamiana* and synthesized by Twist Biosciences with BP-compatible recombination ends. All TIR-encoding ORFs were BP-cloned into the shuttle vector pDONR207 using Gateway^TM^ BP Clonase (Invitrogen) and sequence-verified (GeneWiz). ORFs were then cloned using Gateway^TM^ LR Clonase (Invitrogen) into previously described binary vectors containing Omega leader sequences: pGWB602 (35S Omega:), pGWB615 (35S Omega: HA-), pGWB641 (35S Omega: -GFP), pGWB602 (35S: HA_SAM-) or pGWB602 35S: HA_SAM-intron-). The pGWB602 35S: HA_SAM-intron vector was constructed by adding a *Glycine max* intron from *Glyma18G022500* into the SAM ORF. The polymerase incomplete primer extension (PIPE) method was used to introduce single amino acid substitutions or small insertions/deletions (77). PCR was performed on genes within pDNR207 constructs with Q5 High-Fidelity polymerase (New England Biolabs, Ipswich MA), and all constructs were sequence-verified by GeneWiz prior to cloning into binary vectors. The HA_SAM oligomerization domain (1X HA tag-SAM_478-578_-GGGGS) of SARM1, and the SARM1-TIR domain (561–724) were described previously (25). Amino acid sequences for all of the cloned Arabidopsis TIRs are provided in the Supporting Information.

### Phylogenetic Tree Construction/Multiple Sequence Alignment

*Arabidopsis thaliana* TIR protein sequences were aligned using Muscle within MEGA7 (78). Maximum likelihood trees were then constructed based on full-length sequences.

### Transient Expression in *Nicotiana benthamiana* via *Agrobacterium* Delivery

*Agrobacterium tumefaciens* strain GV3101 was infiltrated using needleless syringes into leaves of ∼4-5 week-old *Nicotiana benthamiana* (*Nb*). GV3101 was infiltrated at OD_600_ of 0.80, except for cell death titration analyses, where the delivered OD is noted within figure panels. The viral suppressor of silencing, p19, was included in the inoculum at OD 0.05, as described previously (25, 48). For EDS1-dependent cell death assays, *Nb* leaves were covered with foil for 2 days and scored for cell death at 5 dpi. GV3101 was transformed via heat shock with ∼500 ng of binary construct DNA. GV3101 liquid cultures were grown overnight at 28°C under selection (50 μg/mL each of rifampicin, gentamicin, spectinomycin or kanamycin, respectively). Cultures were pelleted at 2600 x g for 5 min and then induced for ∼3 h in 10 mM MES buffer (pH 5.6), 10 mM MgCl_2_, and 100 μM acetosyringone prior to leaf infiltration. *N. benthamiana* plants were grown in Conviron or Percival plant growth chambers at 25°C, 70% relative humidity and 15 h photoperiod with a light intensity of ∼80 μE·m^−2^·s^−1^.

### *In planta* LC-MS Analysis

LC-MS/MS analysis was carried out on Waters TQ-XS triple quad mass spectrometer coupled with a Waters Acquity H-class UPLC-C18 column (1.7 μm, 2.1 x 50 mm). Mobile phases were A: 2 mM ammonium acetate, and B: 100% methanol. Flow rate: 0.2 mL/min. Gradient: 0-5 min, 0% B; 5-7 min, 0% to 20% B; 7-8 min, 20% to 100% B; then the column was washed with 100% B for 2 min before equilibration to 100% A over 15 min. Mass spectrometer conditions: Capillary voltage: 800 V; desolvation temperature: 600°C; desolvation gas (nitrogen, 1000 L/h); cone gas: 150 L/h, nebuliser gas: 7 bar. MRM parameters for the detection of cADPR-isomers, 542/136 (cone voltage 20 V, collision energy: 32 eV); 542/348 (cone voltage 20 V, collision energy: 28 eV). cADPR-isomers were verified by comparing with pure standards of 2’cADPR or 2’RFA (2′-O-β-D-ribofuranosyladenosine).

### Immunoblots

Three 6 mm discs from three different *Nb* leaves were harvested into 2.0 mL tubes containing single glass beads and flash frozen in liquid nitrogen. Leaf tissues were homogenized in a TissueLyzer II (Qiagen) at 30 Hz for 30 s. Proteins were extracted in 200 μL of lysis buffer [50 mM Tris·HCl (pH 7.5), 150 mM NaCl, 5 mM EDTA, 0.2% Triton X-100, 10% (vol/vol) glycerol, 1/100 Sigma protease inhibitor cocktail]. Lysates were centrifuged at 4°C for 5 min at 2300g (5K RPM, Beckman FA241.5P rotor) and then stored on ice. Equal lysate volumes were loaded for separation and analysis by SDS-PAGE. Samples were transferred overnight at 4°C in 1X Towbin buffer. Following transfer, nitrocellulose blots were blocked for 1 h at RT (room temperature) in 3% (wt/vol) nonfat dry milk 1X TBS-T (50 mM Tris, 150 mM NaCl, 0.05% Tween 20). Immunoblots for primary antibodies [HA (3F10, Roche) or GFP (Cat# 1181446001, Roche)] were incubated for 1 h at RT in 3% (wt/vol) nonfat dry milk TBS-T at 1:2,000, followed by 3X 5 min washes in 1X TBS-T. Secondary HRP-conjugated goat anti-rabbit or goat anti-mouse IgG (Millipore Sigma) was added at 1:10,000 in TBS-T (3% milk) and incubated for 1 h at room temperature on a platform shaker, followed by 3X 5 min washes in 1X TBS-T. Chemiluminescence detection was performed using Clarity Western ECL Substrate (Bio-Rad) and developed with a ChemiDoc MP chemiluminescent imager (Bio-Rad).

### *In planta* NADase Assays

Two 7 mm discs of transformed *Nb* leaf tissue were cored and harvested into 2.0 mL tubes containing single glass beads and immediately flash-frozen in liquid nitrogen. Leaf tissue was homogenized using a Qiagen TissueLyzer II at 30 Hz for 30 s. Pulverized tissue was resuspended in 400 μL of ice-cold lysis buffer [50 mM Tris·HCl (pH 7.5), 150 mM NaCl, 5 mM EDTA, 0.2% Triton X-100 and 10% (vol/vol) glycerol] and stored on ice. Lysates were pelleted via centrifugation at 4°C for 5 min at 2300g (5K RPM, Beckman FA241.5P rotor) and stored on ice. Total cellular NAD^+^ was detected using the Amplite Fluorimetric NAD^+^-Assay kit (Cat# 15280, AAT Biosciences). 50 μL of clarified supernatant was used in the assay, according to manufacturer’s recommendations. NAD^+^ detection was performed in 96-well black bottom plates (Costar) on an Infinite M Plex (Tecan) plate reader at excitation (420 nm) and emission (480 nm) using i-control 2.0 software and presented as RFU (relative fluorescence units).

### Artificial TIR Design and Consensus Determination

To determine the consensus amino acid sequence of each Arabidopsis TIR group (‘Death’, ‘All’, or ‘Non’), the corresponding TIR protein sequences were aligned using Muscle within MEGA7 and trimmed at the boundaries of the core TIR domain structure (78). Consensus sequences were calculated with Gene-Calc (https://gene-calc.pl/sequences-analysis-tools/consensus-sequence) at a threshold of 0.05. Tie breaks for consensus residues were selected by randomly picking a tied residue. “Random” artificial TIR sequences were generated by randomly selecting an amino acid from the AtTIRome for each position of the core TIR domain. The random TIR script (add pointer to script) selected with replacement (i.e. an individual AtTIR could contribute more than once to a Random TIR) and avoided picking gaps. Artificial TIR ORFs were codon-optimized for expression in *Nb* and synthesized by Twist Biosciences (Twist Biosciences).

### *In vitro* expression constructs

RPS4, RPP1, and SNC1 TIR domain constructs were generated previously and purified as described (24, 29). Col-0 cDNA encoding the TIR domains of AtTN7 (*At1g72900*; residues 2-179), AtTN10 (*At1g72930*; residues 2-176) and AtTN11 (*At1g72940*; residues 2-179) were cloned into pET-28a(+) with cleavable N-terminal 6-His tags. ORFs encoding the artificial consensus TIRs were synthesized by Genscript and cloned into pET-30a(+) with C-terminal 6-His tags.

### Protein purification

pET-28a(+) and pET-30(+) constructs were transformed into BL21 *E. coli* and grown in 1 L of ZYM-5052 auto-induction medium with kanamycin selection at 37°C. Once the OD_600_ reached ∼0.6, the temperature was dropped to 16°C for 20 h. Cells were harvested through centrifugation at 6,000 x g for 25 min and stored at −80 °C until purification. Cell pellets were resuspended in 3X volume of wash buffer (50 mM HEPES, 500 mM NaCl, 30 mM imidazole, 1 mM DTT, pH 7.5) with 1 mM PMSF and 2X EDTA-free protease inhibitor cocktail (Millipore Sigma), lysed by sonication and clarified by centrifugation at 18,000 x g for 50 min at 4 °C. The supernatant was loaded onto a 5 mL HistrapFF column (Cytiva) pre-equilibrated with wash buffer and washed with 10 column volumes of wash buffer. Protein was eluted using 50 mM HEPES (pH 7.5), 500 mM NaCl, 300 mM imidazole, and 1 mM DTT. Eluted proteins were then 2X diluted with gel filtration buffer (GFB; 10 mM HEPES, 150 mM NaCl, 1 mM DTT, pH 7.5). For ‘All’ and ‘All’ + ‘Death’ Loop TIR domains, a high-salt GFB was used (20 mM HEPES, 500 mM NaCl, pH7.5). Sample (10 mL) was injected onto a Superdex S75 pg 26/600 size exclusion column (SEC) and fractions were analyzed using SDS-PAGE. Samples containing protein of interest were pooled, concentrated to a minimum of 5 mg/mL, and stored at −80 °C.

### *In vitro* NADase kinetic assays

100 µM of each TIR domain was incubated with 2 mM NAD^+^, 2 mM ATP, and 15 µL of Ni-NTA beads in a total reaction volume of 90 µL in GFB (see above) at 25°C. Time points were taken at 0 h, 0.5 h, 1 h, 1.5 h, 4 h, 22 h and 45 h. To remove Ni-NTA beads, samples were centrifuged for 20 s at 5,000 x g at each time point, and 10 µL of supernatant was transferred to a new tube. Reactions were quenched prior to NMR measurements, by being diluted 5X in GFB and transferred to 150 µL of NMR quench mixture (200 µL final volume with 20 µM DSS (sodium trimethylsilylpropanesulfonate), 1% HCl, and 5% D_2_O). To remove any precipitant, samples were centrifuged at 12,000 x *g* for 5 min and the supernatant was transferred into 3 mm Bruker sample jet tubes. Spectra were recorded using a Bruker Neo 900 MHz NMR spectrometer equipped with a cryogenically cooled probe. Water suppression was achieved using excitation sculpting (180° water-selective) and a flip back pulse (zgesfpgp). The spectra were acquired using 512 scans with an interscan delay of 1 s, at 298 K. All spectra were processed and analyzed using TopSpin™ (Bruker). NAD^+^ cleavage was validated by detecting Nam. NAD^+^ was quantified as a relative % for each time point, compared to 0 h NAD^+^ intensity, and calculated in triplicates. The NAD^+^ peak intensity (9.4 ppm) was fit to a Lorentz-Gauss function (Voigt) within Topspin). All spectra were referenced to DSS.

### End point product assays

These assays were performed as the kinetic NADase assays except that 10 µL (RPS4-TIR), 25 µL (RPP1-TIR, TN7-TIR, RPP1-TIR, TN10-TIR) or 40 µL (SNC1-TIR, TN11-TIR and consensus TIRs) of reactions were diluted to 50 µL and transferred to the NMR quench mixture. For MALDI-TOF-MS (matrix assisted laser deionization – time of flight mass spectrometry), 5 µL of the reaction mixture was diluted with 5 µL of ddH_2_O. For TIR product determination, reactions were compared to standards of cADPR, 2’cADPR, 3’cADPR, pRib-AMP, ADPR, ATP, NAD^+^, or Nam at 50 - 75 µM in reaction buffer.

### MALDI-TOF MS

NADase reactions were boiled at 90°C for 5 min, centrifuged at 10,000 x g and transferred to a new tube. A supersaturated solution of α-cyano-4-hydroxycinnamic acid (CHCA) in acetonitrile was diluted 10X in ethanol: acetone: 0.1% trifluoroacetic acid (TFA) (6:3:1) solution and used as a matrix. 2 µL matrix was mixed with 1 µL sample. Each sample was spotted twice (∼1.5 µL each) on a MTP 384 ground steel MALDI plate. MS was performed on a Bruker TIMS TOF Flex - MALDI-2 with TIMS TOF Control 5.1, a calibration was performed using ESI calibration solution (Fluka 63606), where error was less than 0.5 ppm for all samples. Laser power was set to 62% at a repetition rate of 1000 Hz. An MS scan was performed in positive QqTOF mode over a range of 200 - 2000 m/z. MS parameters were tuned to detect compounds from 200 – 1500 m/z using funnel 1 RF: 350 Vpp; isCID: 0.0 eV; funnel 2 RF: 350 Vpp; multipole RF: 350 Vpp; collision energy: 10 eV; collision RF: 1500 Vpp; quadrupole settings: ion energy: 5 eV; low mass: 250 m/z; focus pre TOF settings: transfer time: 80 µs; pre pulse storage: 10 µs; and detection was set to focus mode. MS/MS fragmentation was performed with a width range 0.8 *m/z* and 15 eV collision energy, and summed with second round collisions at 30 eV. MS tune parameters were adjusted for <200 m/z for 462.1 m/z compound fragmentation as follows; funnel 1: RF 300 Vpp; isCID: 0.0 eV, funnel 2: RF 200 Vpp; multipole RF: 200 Vpp; collision energy: 10 eV; collision RF: 450 Vpp; quadrupole settings: ion energy: 5 eV; low mass: 60 m/z; focus pre TOF settings: transfer time: 65 µs; pre pulse storage: 3 µs.

### Protein Structure Modeling

Protein structure models for artificial TIR proteins, ‘All’, ‘Death’ and ‘Non’ were generated using AlphaFold3 (79). The resulting PDB files were analyzed and labeled using PyMOL (The PyMOL Molecular Graphics System, Version 1.8 Schrödinger, LLC).

### Statistical Analyses

Multiple comparisons were analyzed via one-way ANOVA with post-hoc Tukey HSD (honestly significant differences) using R Studio (version 1.4.1717, RStudio Team (2021), Boston, MA). Comparisons of significance are indicated with compact letter display (CLD). Over-lapping letters are non-significant (*p*>0.05), while separate letter classes indicate *p*<0.05 or better.

## Supporting information

Supplemental Script

## Acknowledgements

We would like to thank Sarah Grant, Paulo Teixeira, and Farid El-Kasmi for careful reading and discussion of the manuscript. We also thank David Cook, Emily Hudson-Arns, Eliza Walthers, Christopher Gomez, Gustavo Contreras, Zac Newland-Smith, and Jaden Eisenach for laboratory assistance. We thank Brett Hamilton and the Centre for Microscopy and Microanalysis for providing support with the use of the TIMs TOF Flex.

## Author contributions

Conceptualization: AMB, LS, MS, JT, NM, QL, JLD, LW, BK, MG, MTN. Methodology & analyses: AMB, LS, MS, AF, JT, SCO, TST, MD, VN, MG, MTN. Writing-original draft: AMB, LS, MG, MTN. Writing Review & Editing: all authors.

## Data and material availability

All data and analyses needed to evaluate the conclusions in the paper are presented in the paper and/or the Supplementary Materials

## SI Figure Legends

**SI 1.**
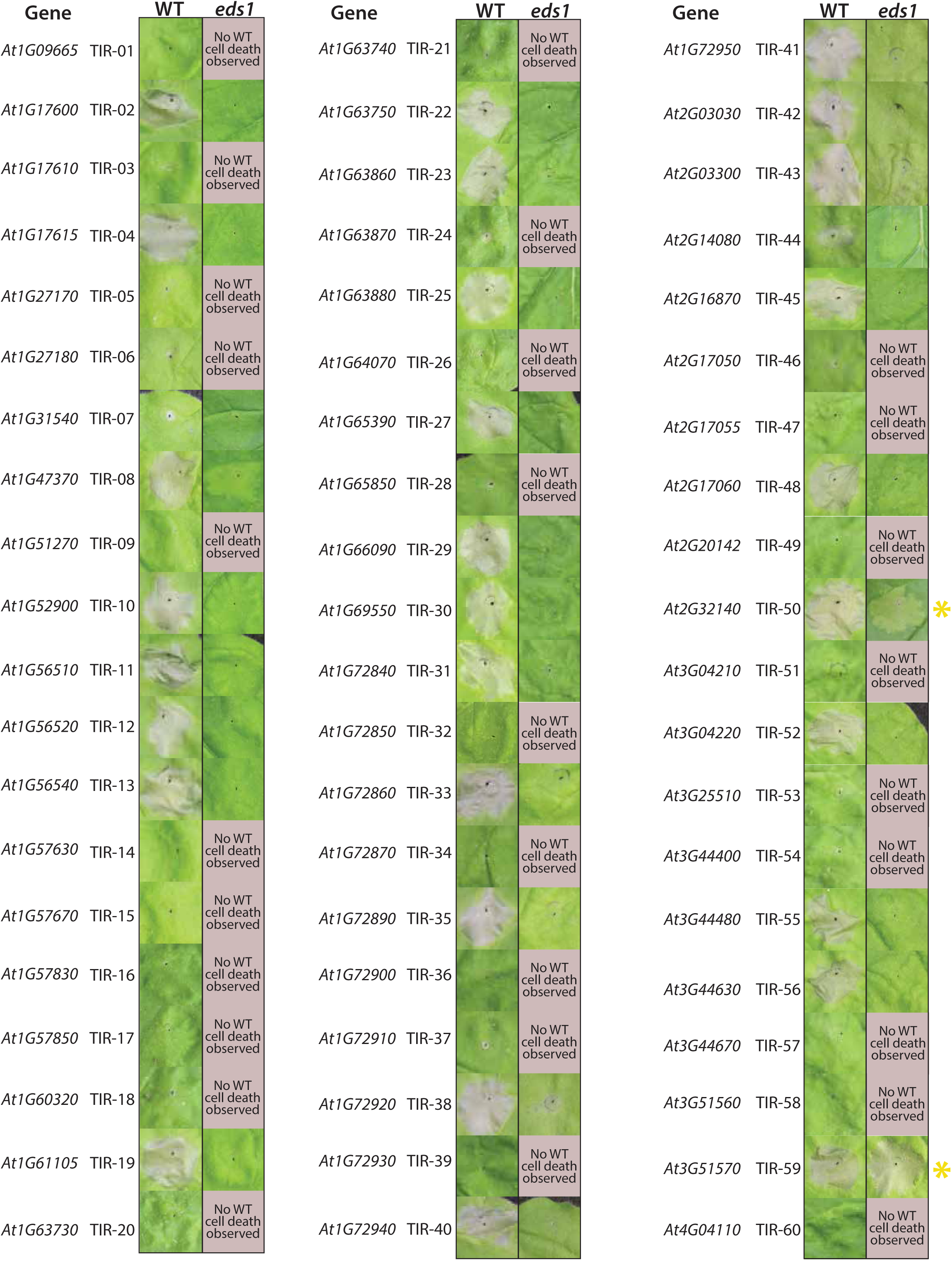
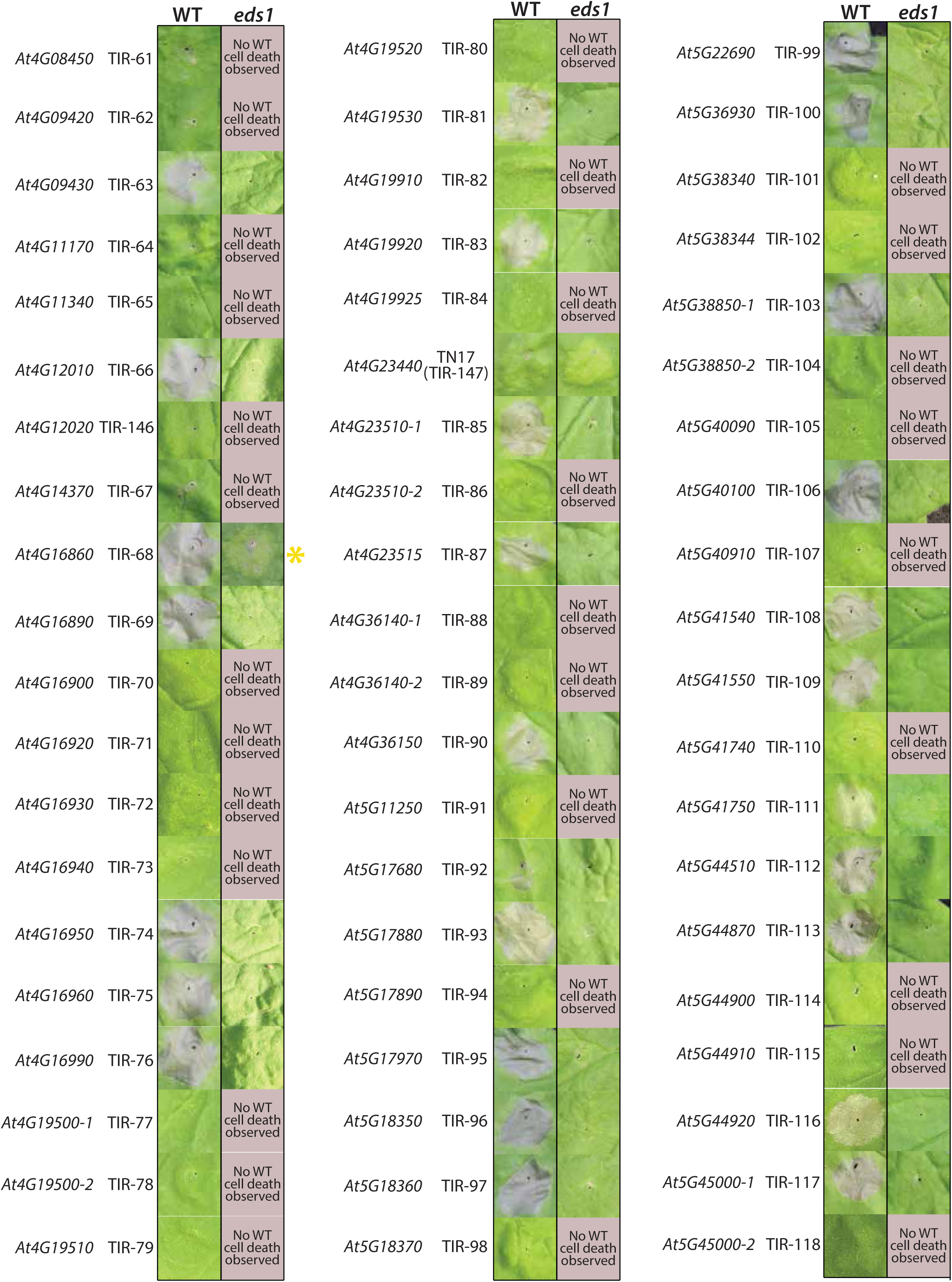
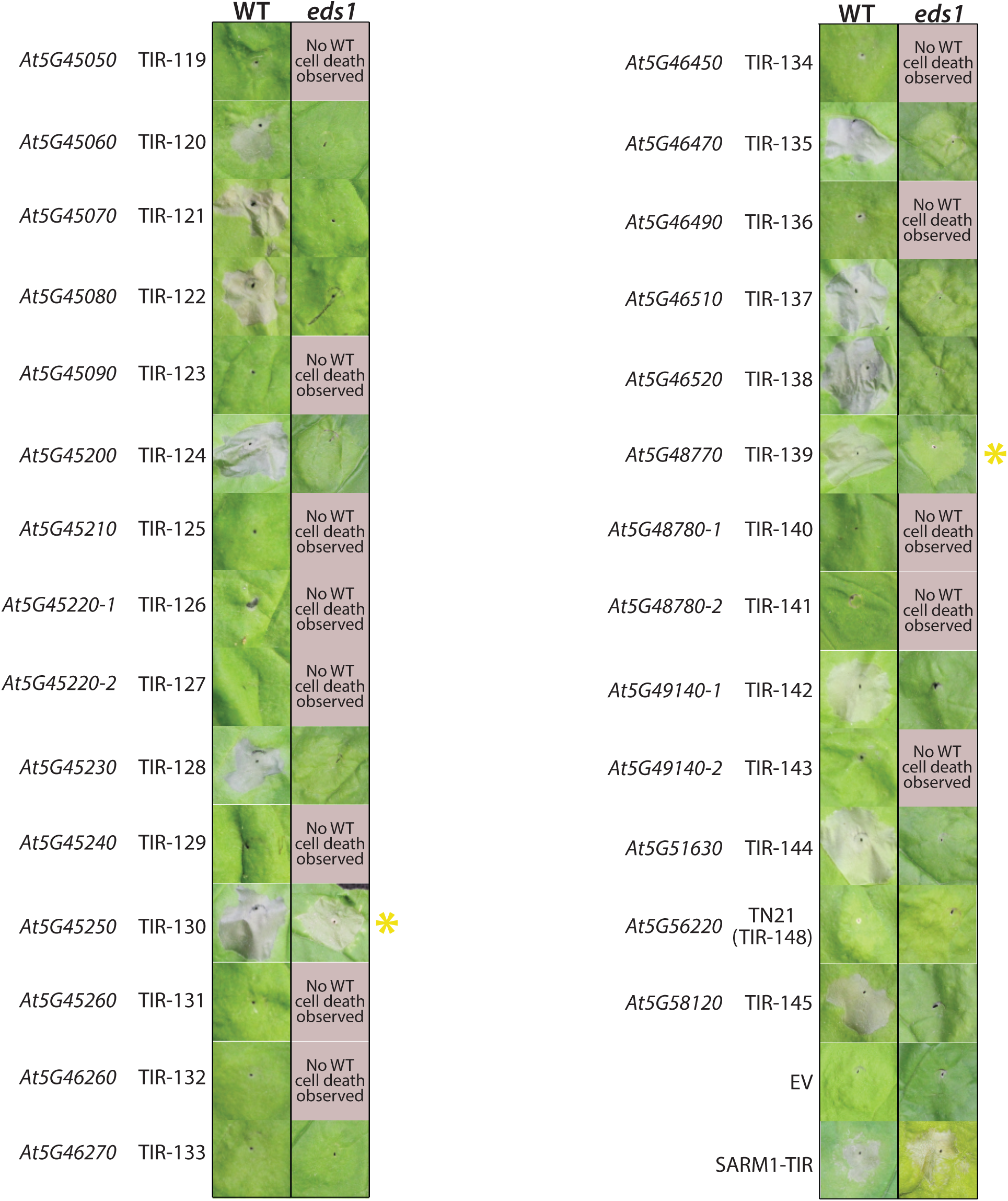
Cell death induction by the Arabidopsis Col-0 TIR-ome. *Nicotiana benthamiana* (*Nb*) WT or *eds1* leaves expressing 35S: HA_SAM tagged AtTIRs. AtTIRs were expressed at OD 0.80 and imaged ∼5 dpi. The WT leaves were kept dark with foil coverings for 48 h after infiltration to enhance *EDS1*-cell death, as reported by Qi *et al* (80). EV (empty vector, 35S: YFP), SAM_SARM1-TIR (sterile alpha and TIR motif containing 1). Similar experiments were performed at least three times. Any AtTIR that triggered cell death in WT was examined for EDS1-independent effects. A yellow asterisk indicates visible induction of minor to moderate chlorosis in *Nb eds1*.

**SI 2.**
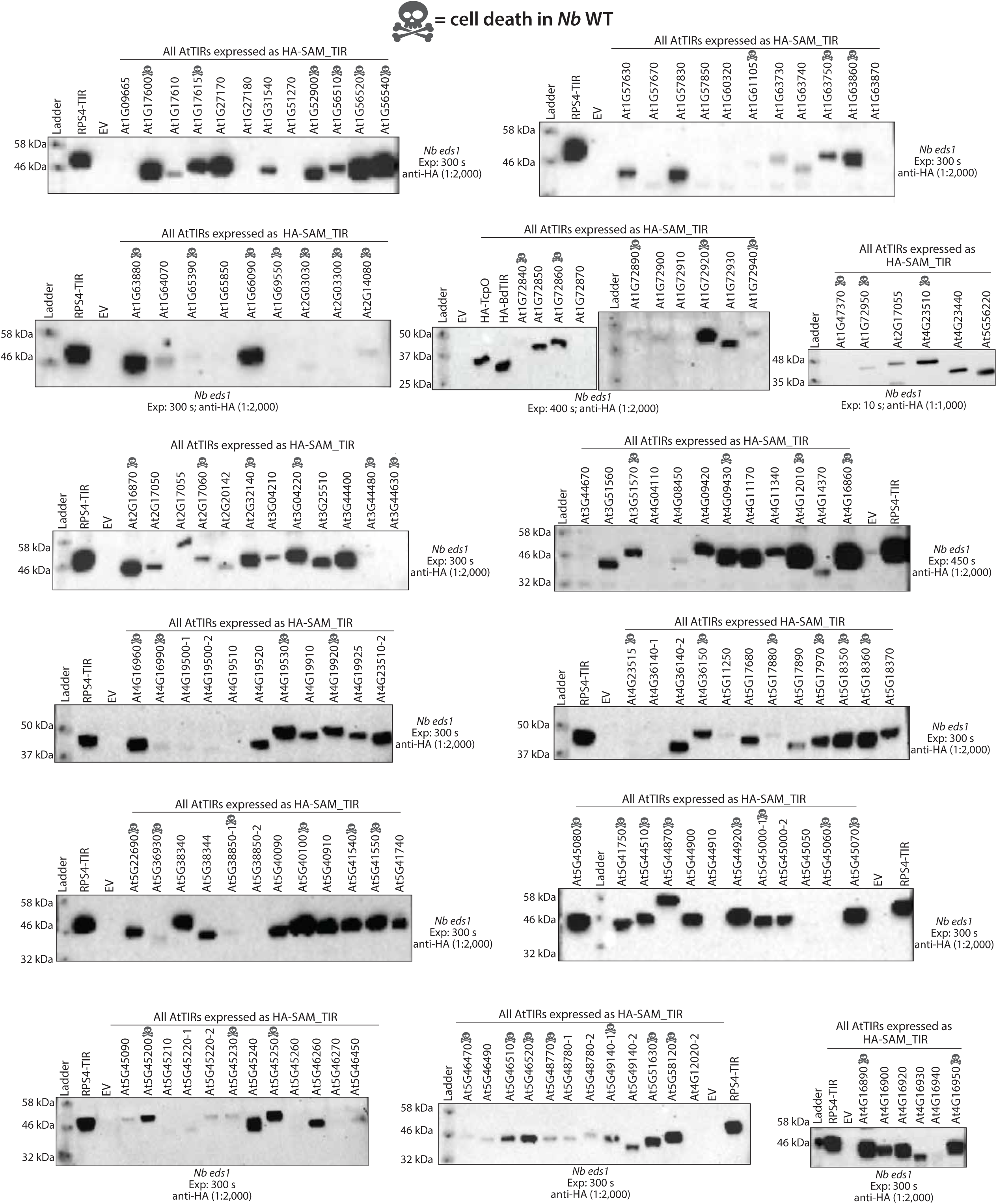
Protein accumulation of the Arabidopsis Col-0 TIR-ome. Anti-HA immunoblot of HA_SAM tagged AtTIRs. AtTIRs were transiently expressed in *Nb eds1* leaves at OD_600_ 0.80. 3X 6 mm leaf discs were collected and pooled at ∼40 hpi for immunoblot analysis. The anti-HA antibody used at 1:2,000 and blots were typically exposed ∼300 s. A skull and crossbones indicates cell death induction in *Nb* WT.

**SI 3.**
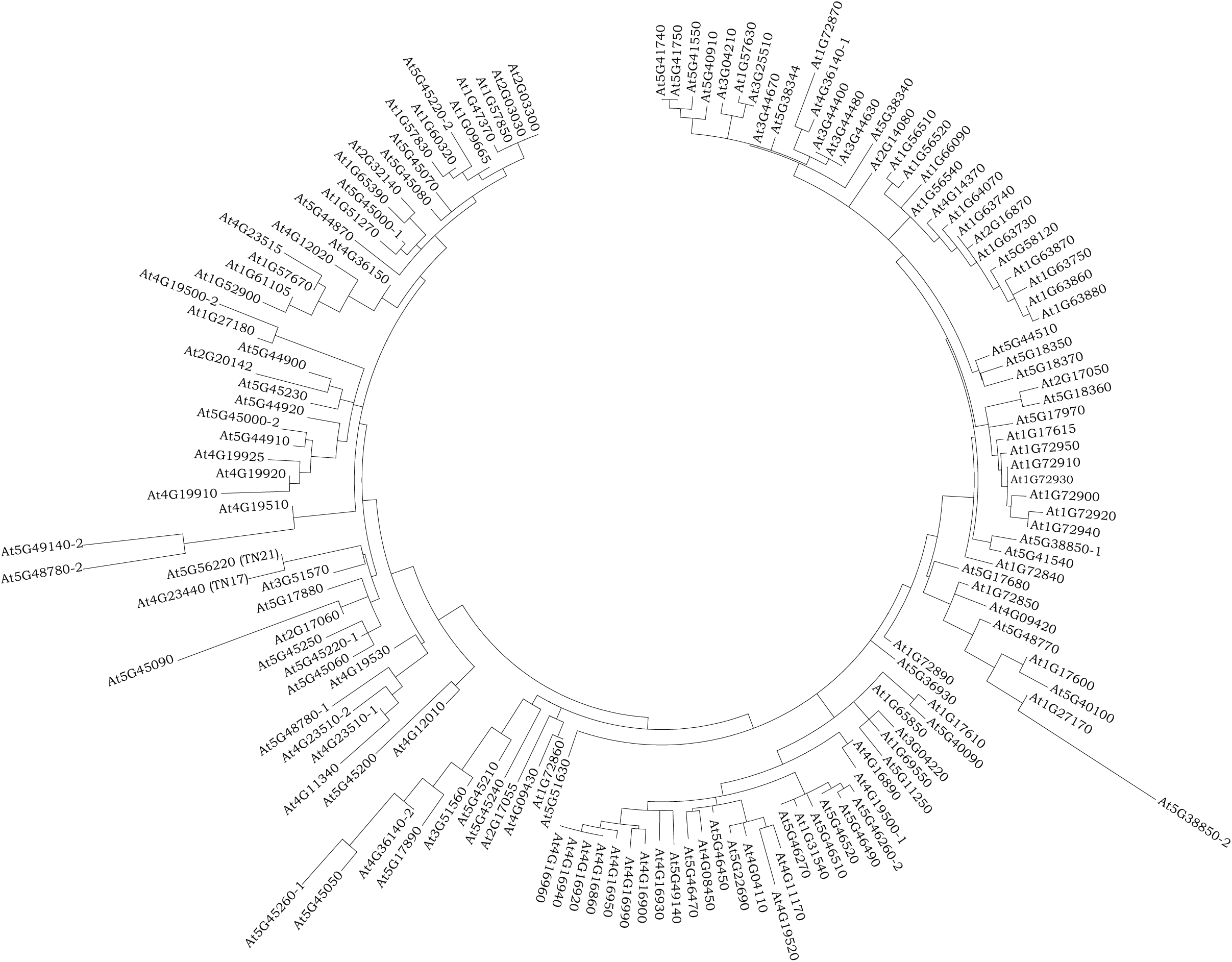
Maximum-likelihood phylogenetic tree of the Arabidopsis thaliana Col-0 TIR-ome. Evolutionary history of expressed Arabidopsis TIRs, as inferred using the maximum likelihood method based on the JTT matrix-based model (Jones et al, 1992). TIR sequences were not trimmed to the minimal core TIR domain and thus include the proceeding or trailing (+80) sequences (Swiderski et al, 2009). The tree with the highest log likelihood (−46771.30) is shown. Initial tree(s) for the heuristic search were obtained automatically by applying neighbor-join and BioNJ algorithms to a matrix of pairwise distances estimated using a JTT model, and then selecting the topology with superior log likelihood value. The analysis involved 148 amino acid sequences and was conducted with MEGA7 (Kumar et al, 2016).

**SI 4.**
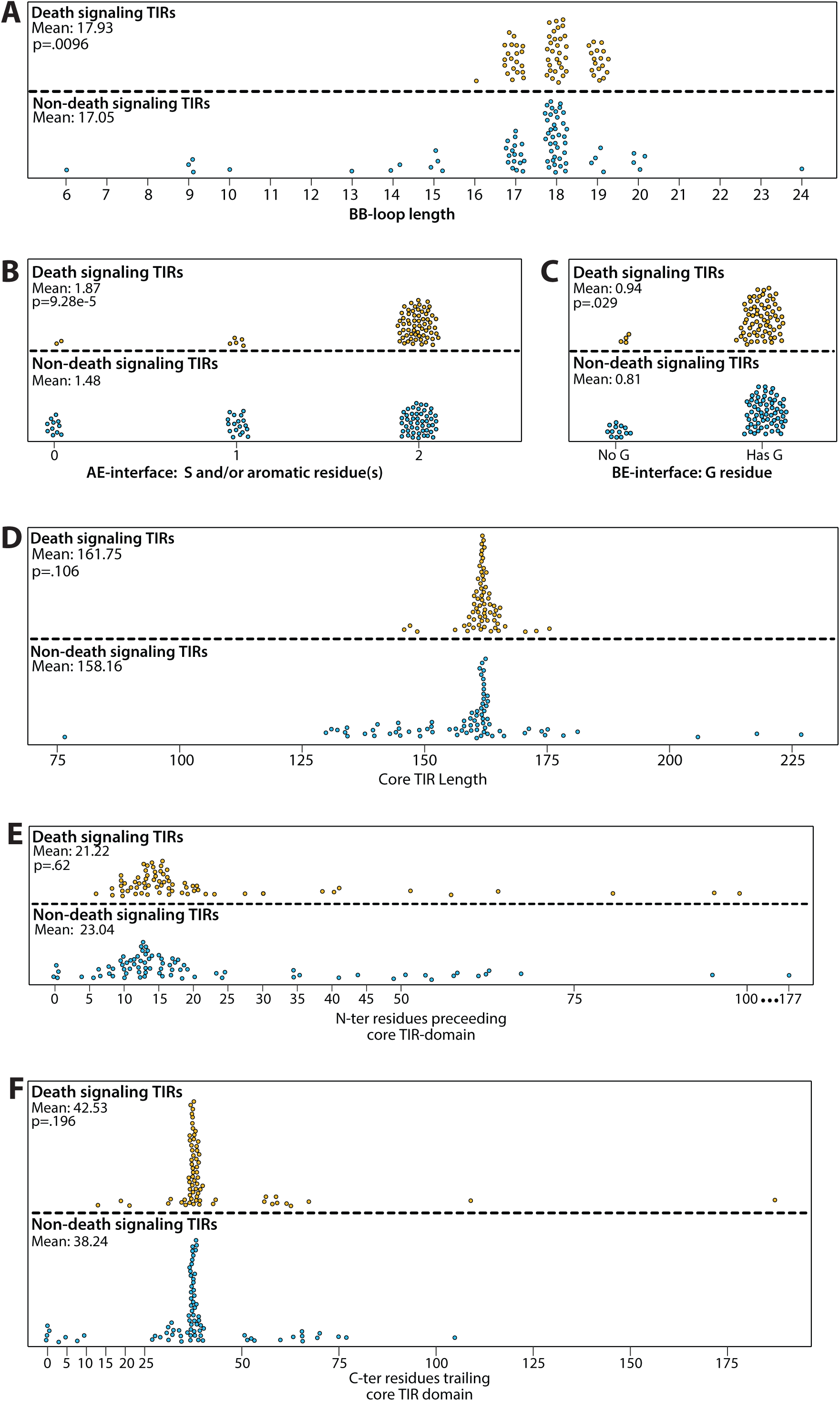
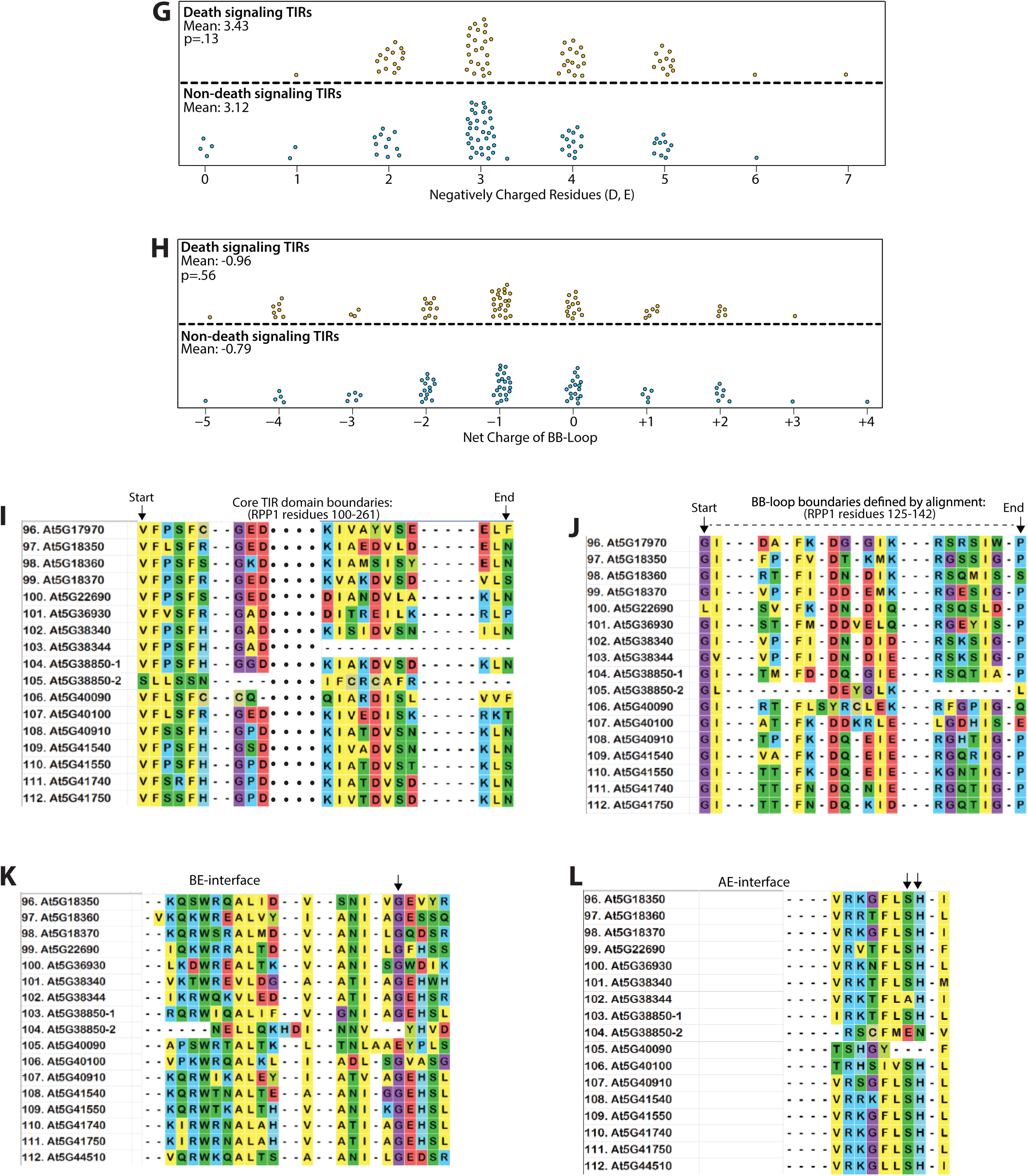
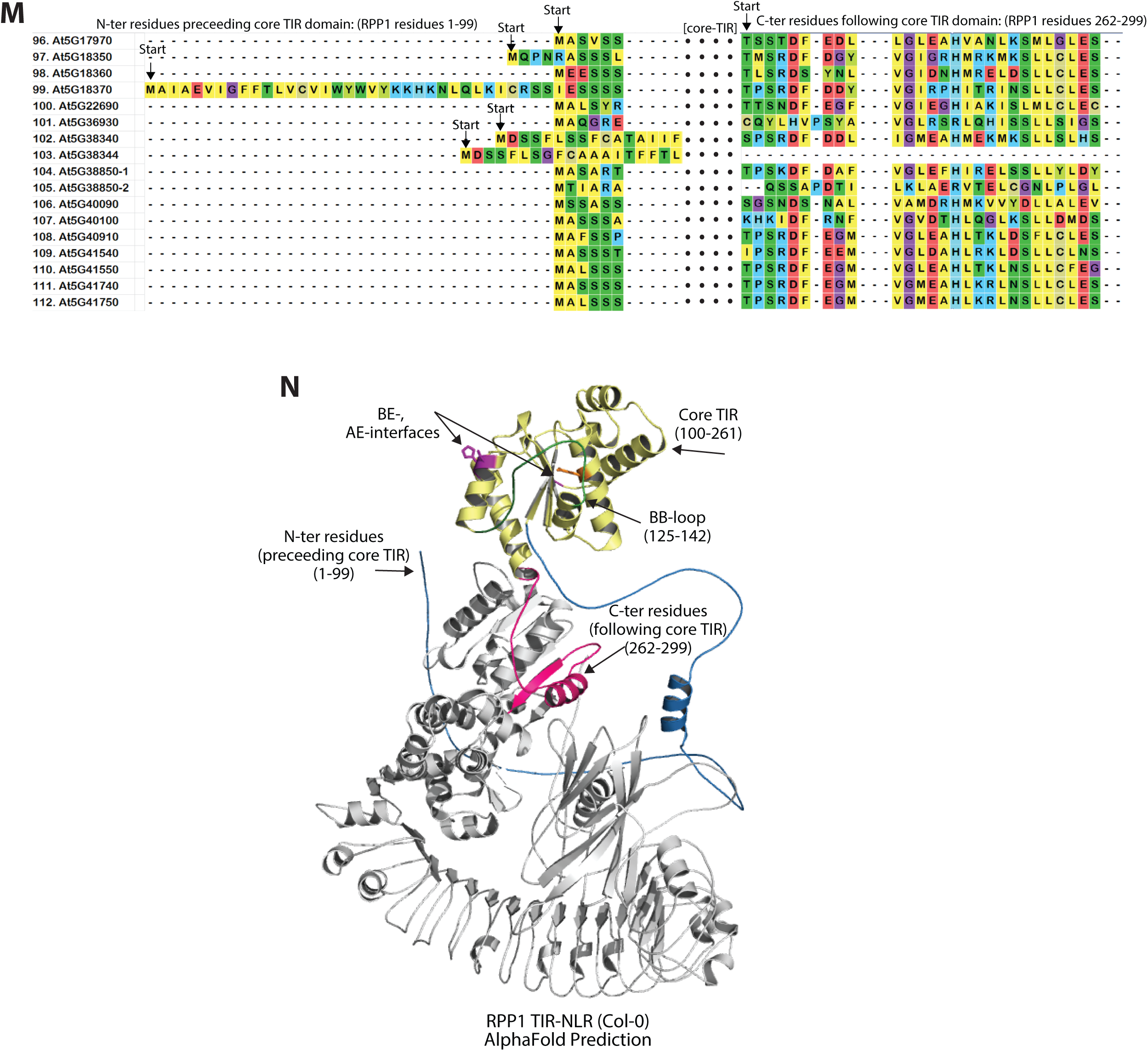
AtTIR-ome *domain metrics*. (*A*) Scatterplot of BB-loop lengths among ‘Death’ and ‘Non’ signaling TIRs. BB-loop boundaries determined by MSA and structural alignment. (*B*, *C*) Scatterplot showing presence/absence of conserved TIR-TIR interface residues (‘AE’ and ‘BE’ interfaces). Canonical ‘AE’ residues: serine or an aromatic residue, histidine (or other aromatic). ‘BE’ residue: glycine (G) (Zhang et al., 2017; Ma et al., 2020; Martin et al., 2020). (*D*-*F*) Scatterplots of minimal core TIR domain length (number of residues), or of N or C-terminal lengths outside the core domain for the ‘Death’ vs. ‘Non’ AtTIR groups. Specific TIR regions were defined by a Muscle alignment of the AtTIR-ome and structural modeling to the RPP1 TIR cryo-EM structure (PDB 7CRC) (Ma et al., 2020). (*G*, *H*) Scatterplots of BB-loop net charge or negatively charged residue abundance among ‘Death’ vs. ‘Non’ signaling AtTIRs. (*I - L*) Muscle alignments (all 148 Col-0 TIRs) used to define TIR domain regions/boundaries (minimal core TIR domain, BB-loop, interfaces). All alignments performed with MEGA7 (Kumar et al, 2016). (*M*) AlphaFold2 model of RPP1 (Col-0) showing the TIR domain boundaries used to calculate the above TIR metrics. The minimal core TIR domain is shown in yellow, BB-loop in green, N-terminal residues preceding the core TIR domain in blue, trailing C-terminal residues in magenta (see arrows), and remaining NBS-LRR region shown in grey. The length of each delineated region is listed on the structure.

**SI 5.**
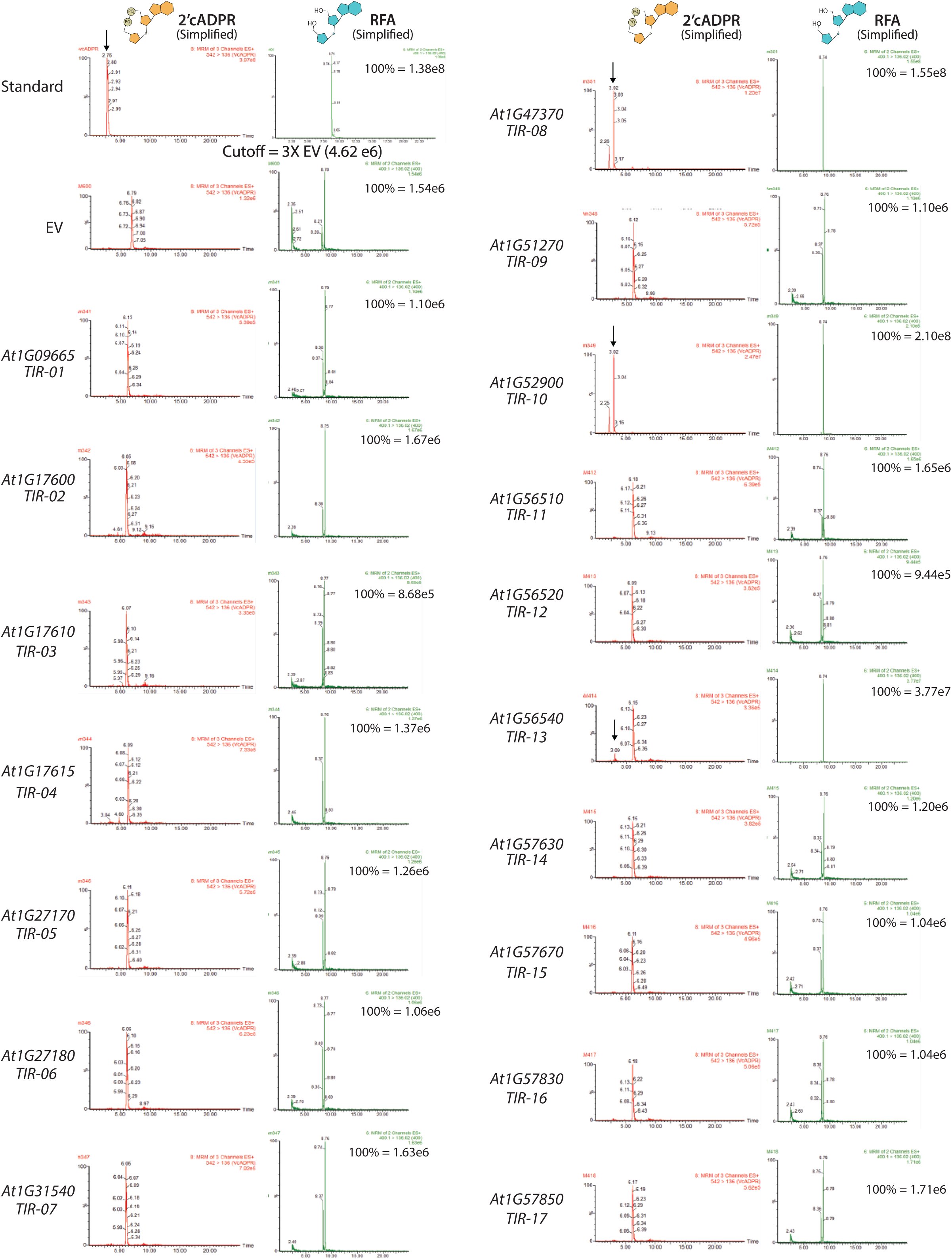
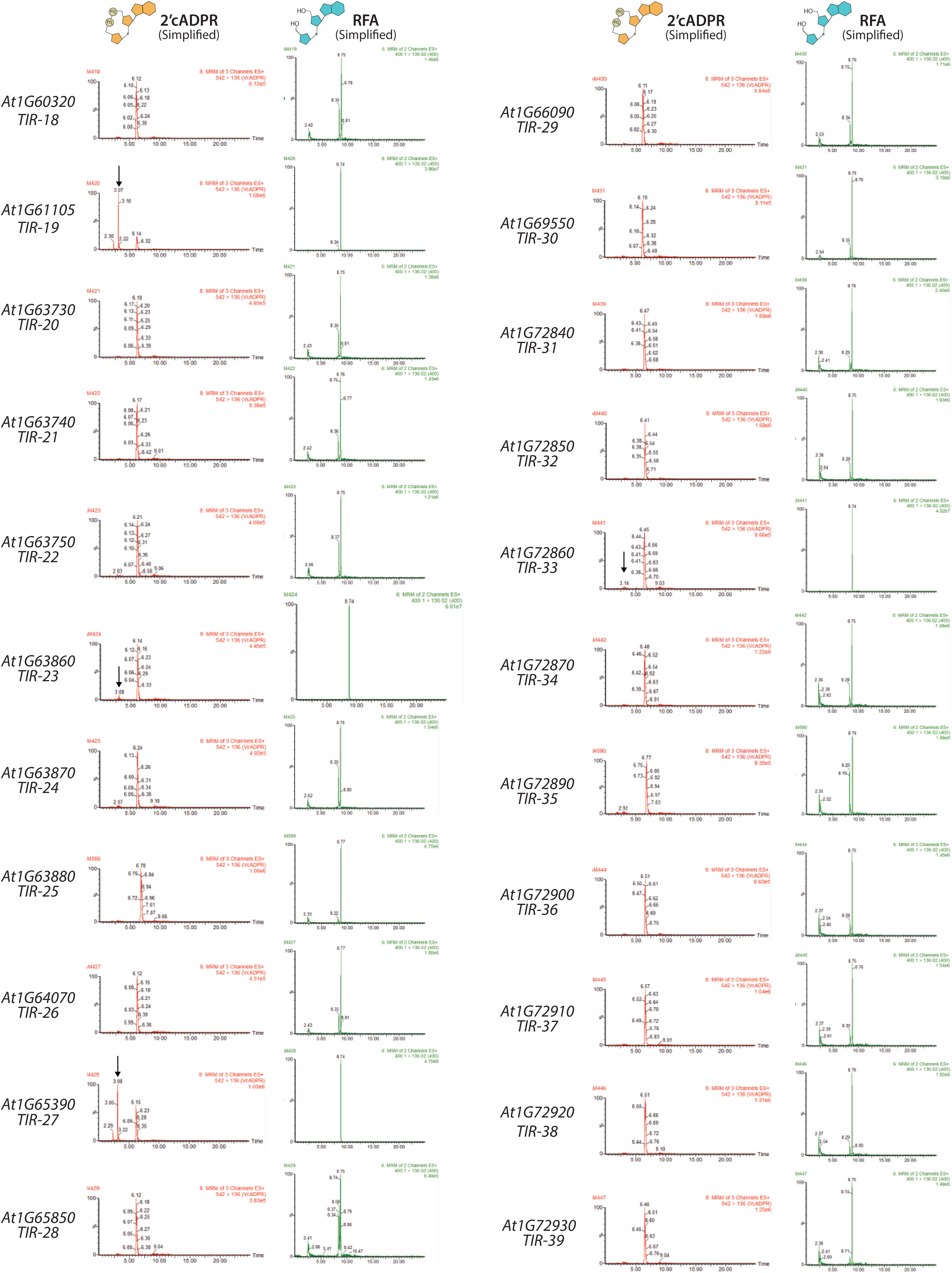
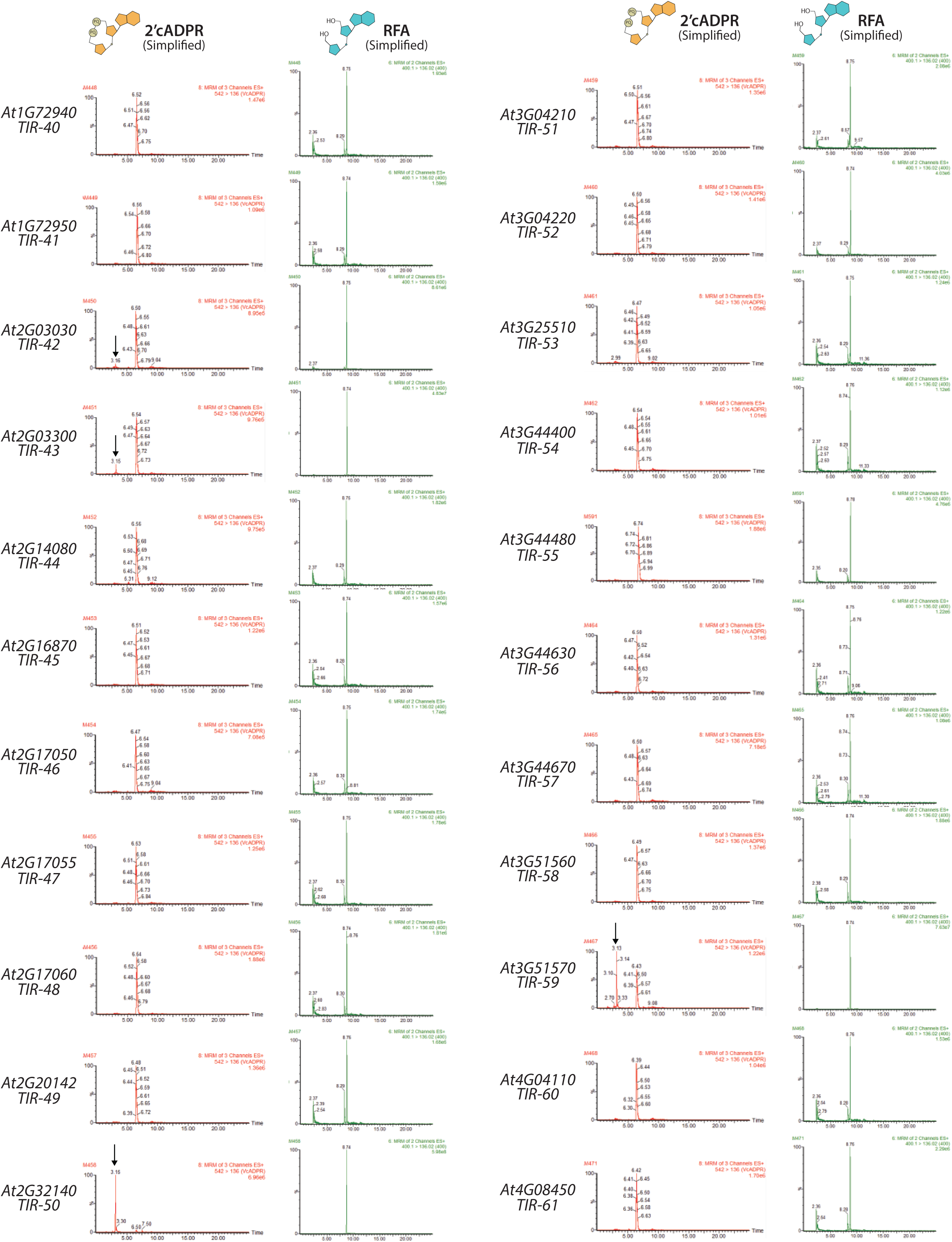
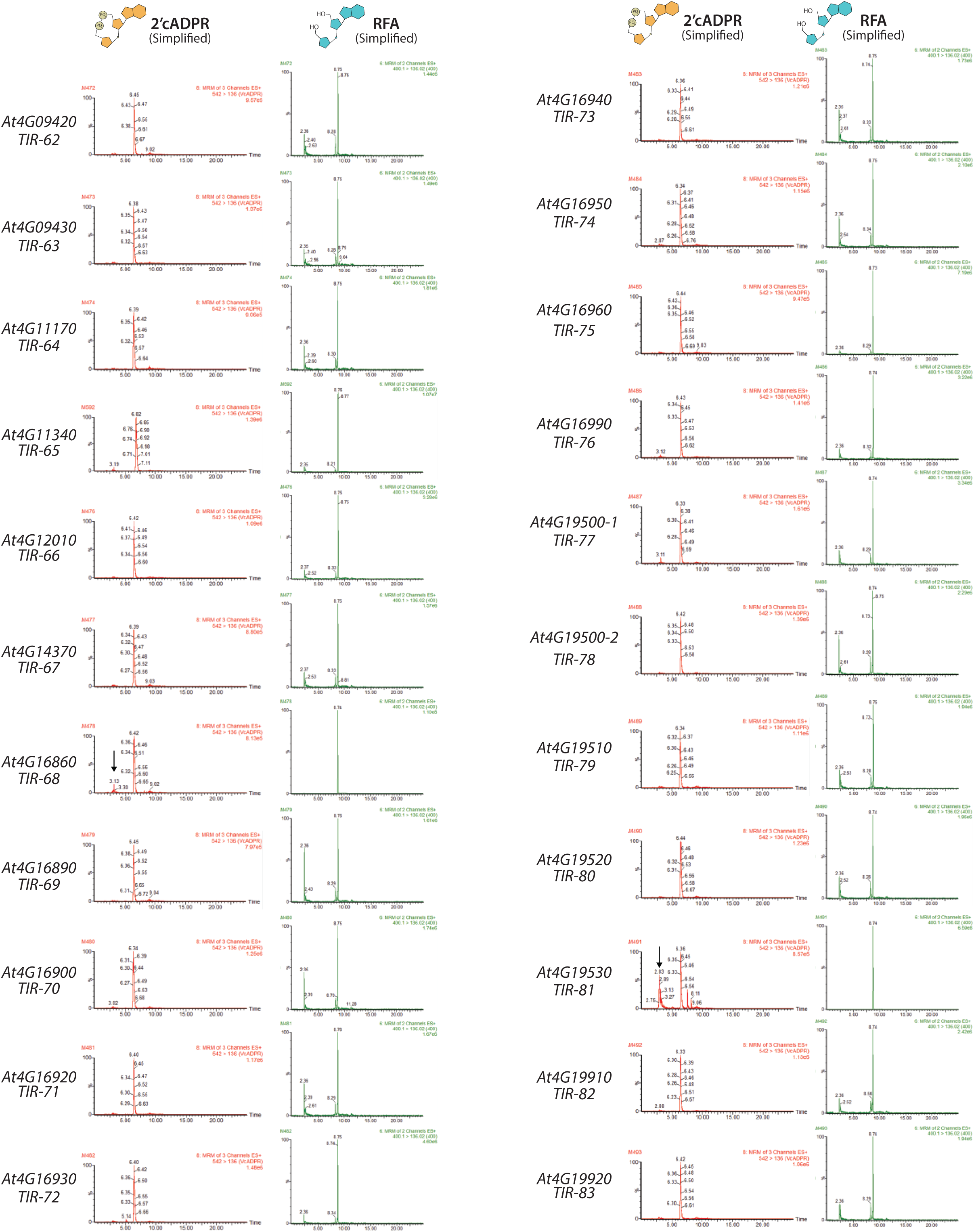
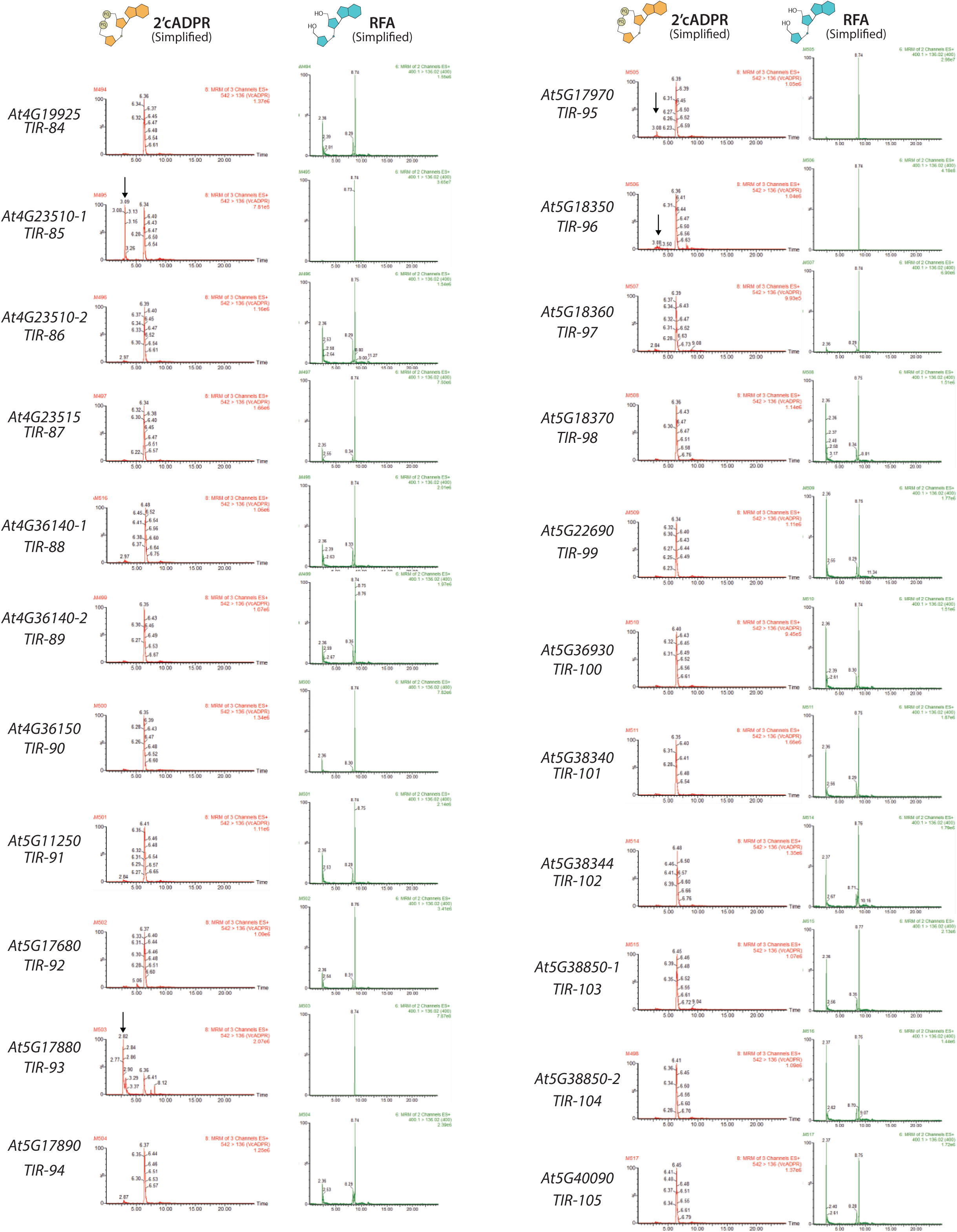
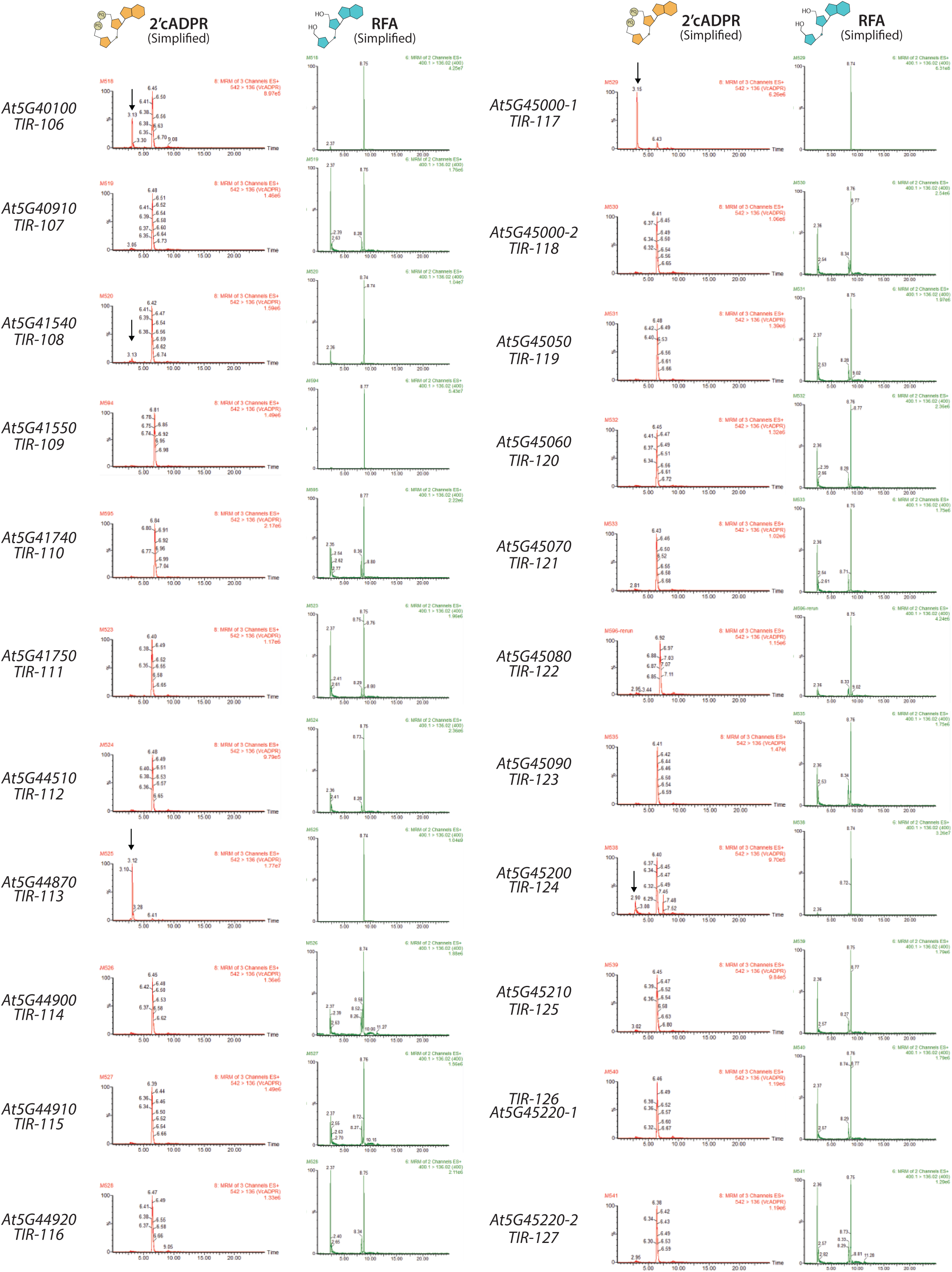
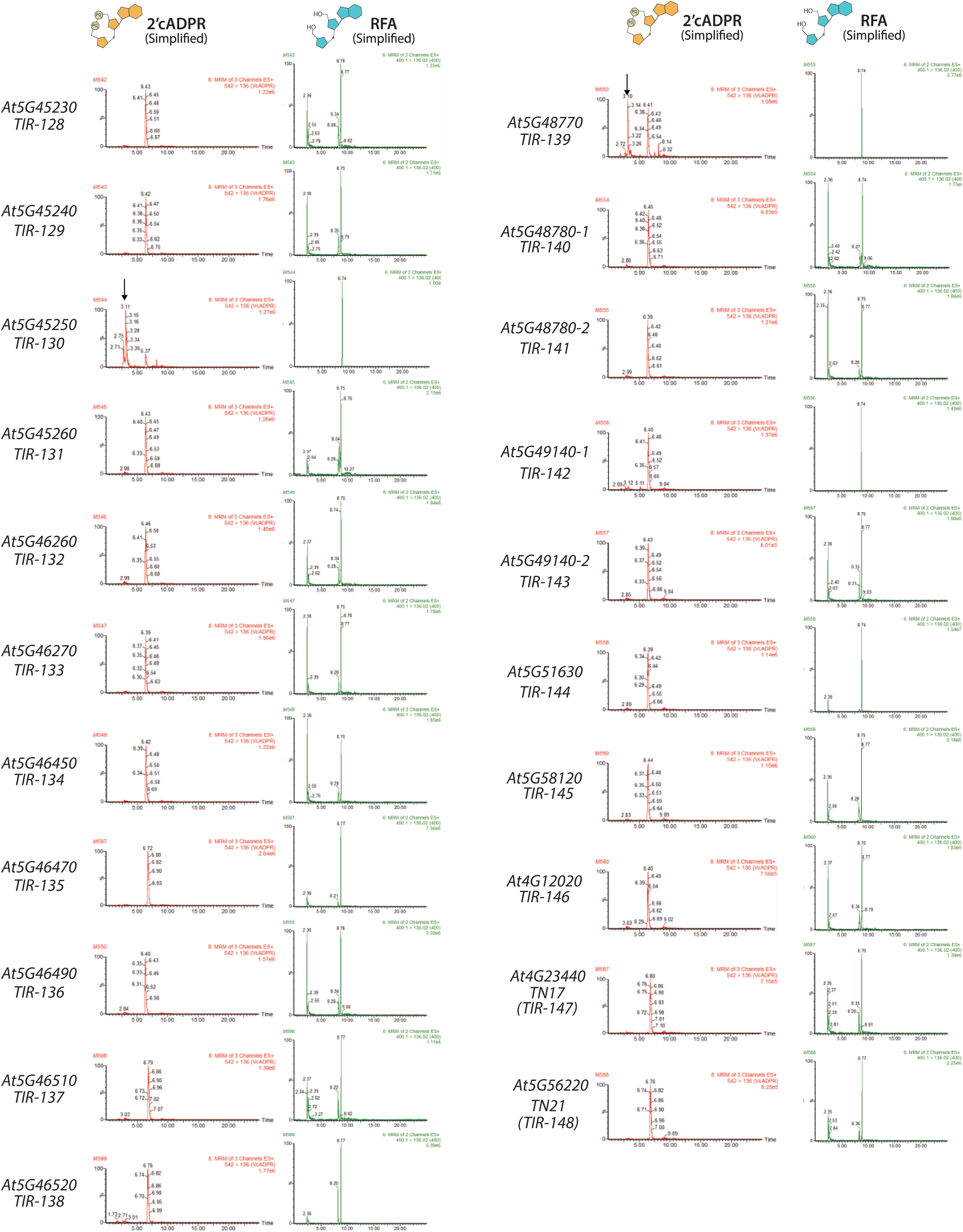
2’cADPR and RFA metabolite production by the Arabidopsis Col-0 TIR-ome. LC-MS chromatograph traces of 2’cADPR (cyclic ADP-ribose) and RFA (ribofuranosyladenosine) from *Nb eds1* leaves expressing HA_SAM AtTIR constructs. EV: empty vector (35S:YFP). Standards of 2’cADPR and RFA were run to verify metabolite retention times. All constructs were infiltrated at OD_600_ 0.80; *Nb eds1* leaves were sampled at ∼40 hpi.

**SI 6.**
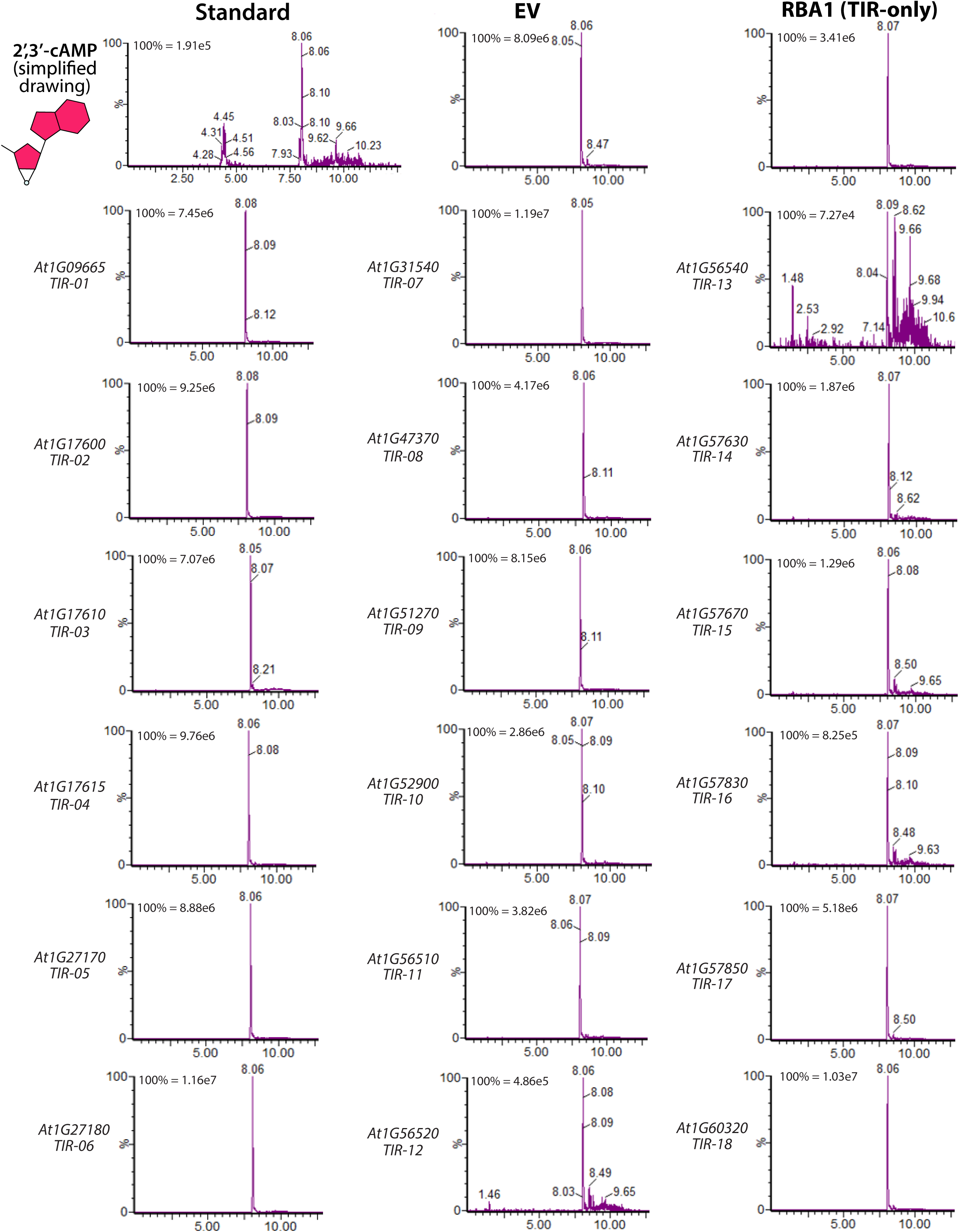
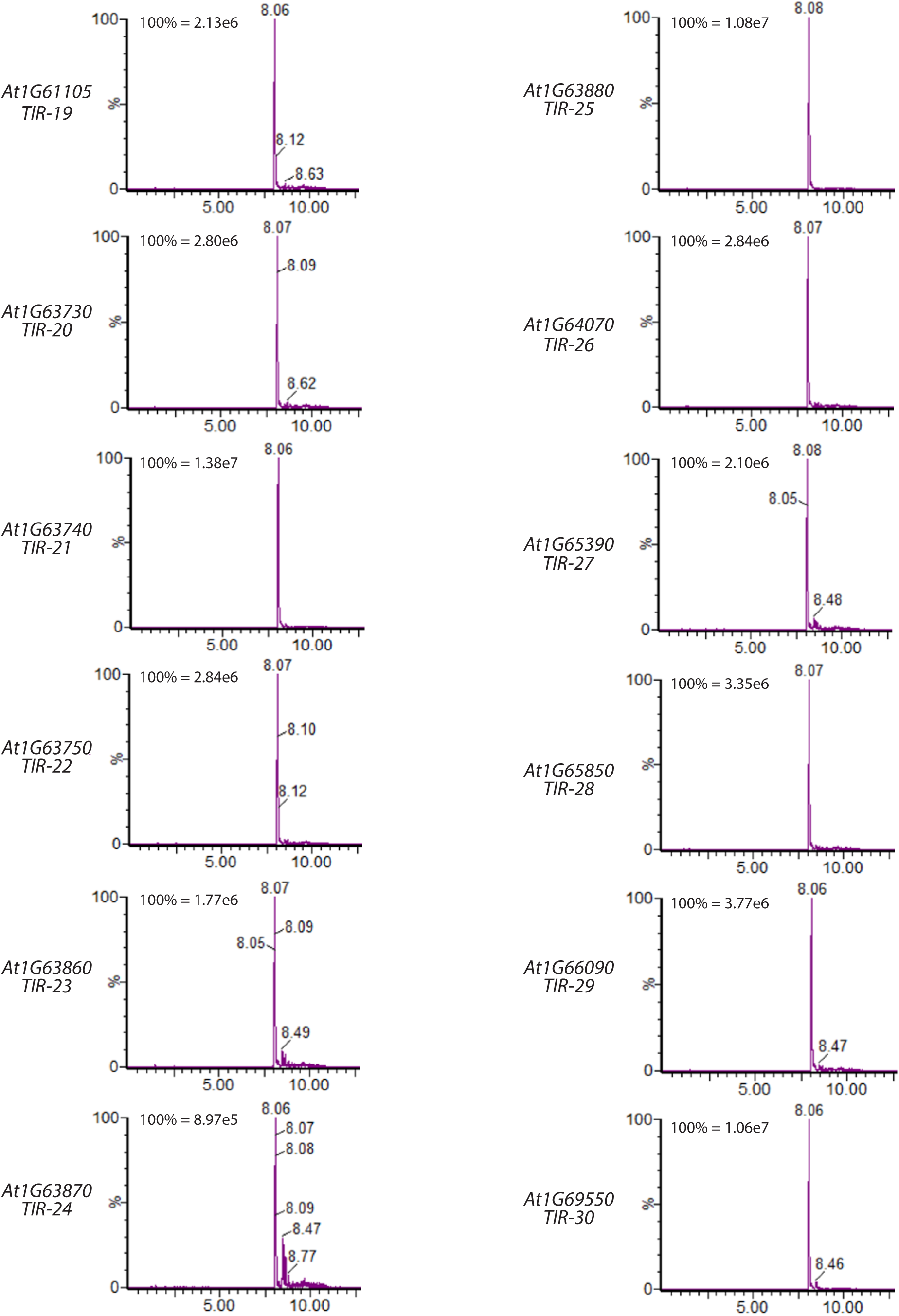
2’,3’-cAMP production by the Arabidopsis Col-0 TIR-ome. LC-MS chromatograph traces of 2’,3’-cAMP in *Nb eds1* leaves expressing HA_SAM AtTIR constructs. Pure 2’,3’-cAMP standard was also run to verify 2’,3’-cAMP retention times. All constructs were infiltrated at OD_600_ 0.80; *Nb eds1* leaves were sampled ∼40 hpi. RBA1:Response to HopBA1; EV: empty vector (35S:YFP).

**SI 7.**
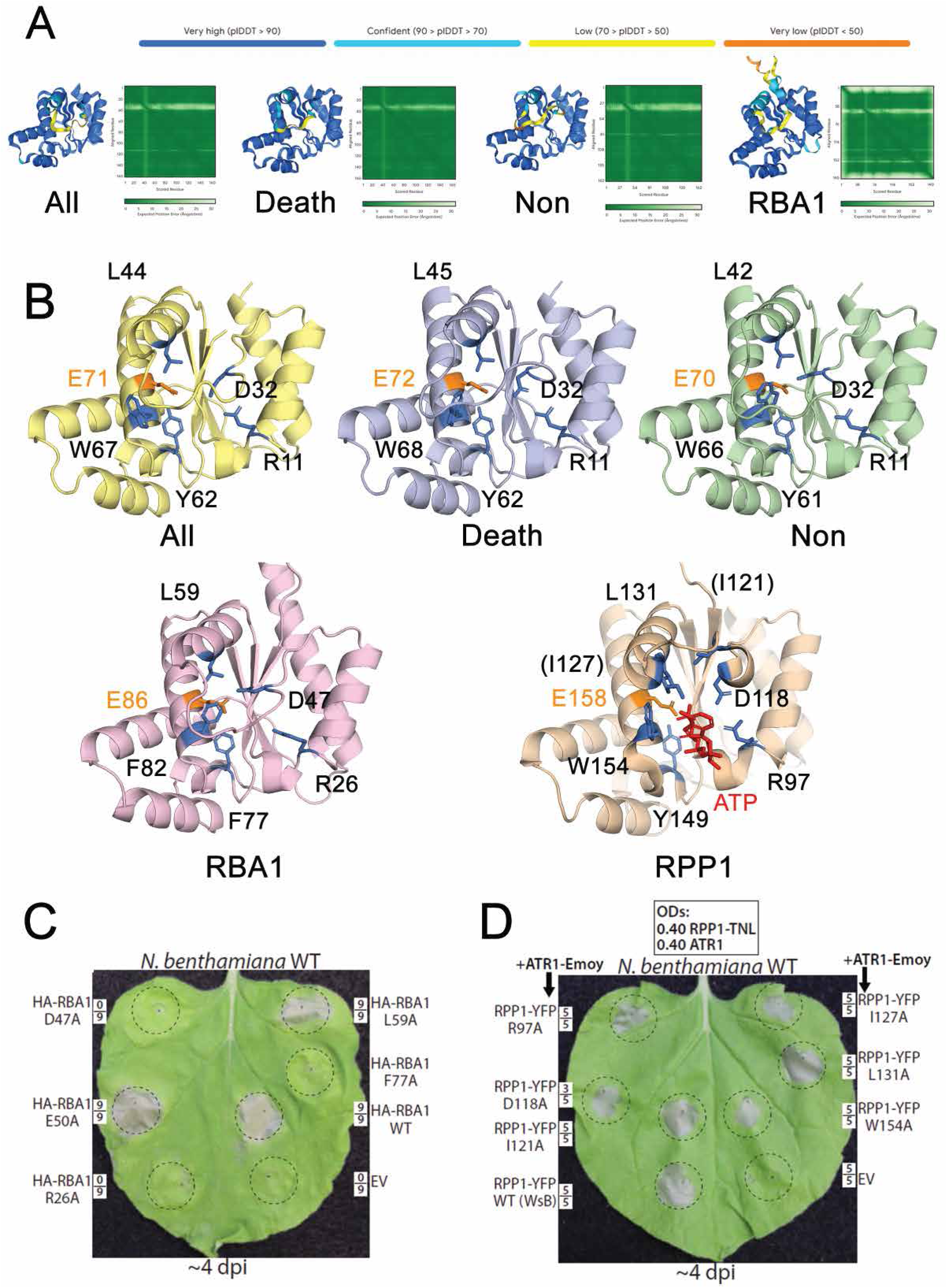
Artificial consensus TIRs have predicted catalytic pockets similar to RBA1 and RPP1; not all proposed catalytic pocket residues are required for signaling. (A) AlphaFold3 models and confidence scores for RBA1 and consensus TIRs (All, Death, and Non) (79). (B) Consensus TIR and RBA1 AlphaFold3 models and RPP1 TIR (PDB 7CRC) labeled to show putative catalytic pocket residues proposed in Manik et al. 2022. Pocket residues labeled in blue; catalytic glutamate labeled in orange. While most residues are well-conserved across TIRs, RPP1 I121 and I127 are not well conserved and are shown in parentheses for clarity. (C, D) N. benthamiana WT leaves expressing HA-RBA1 WT, RPP1-YFP (WsB)/ATR1 (Emoy), or respective catalytic pocket mutant. HA-RBA1 and EV (35S:YFP) constructs were expressed at OD 0.80, while ATR1-Emoy and RPP1-YFP (WsB) were expressed at OD 0.40. Leaves imaged ∼4 dpi and framed numbers denote cell death-positive replicates/total tested.

**SI 8.**
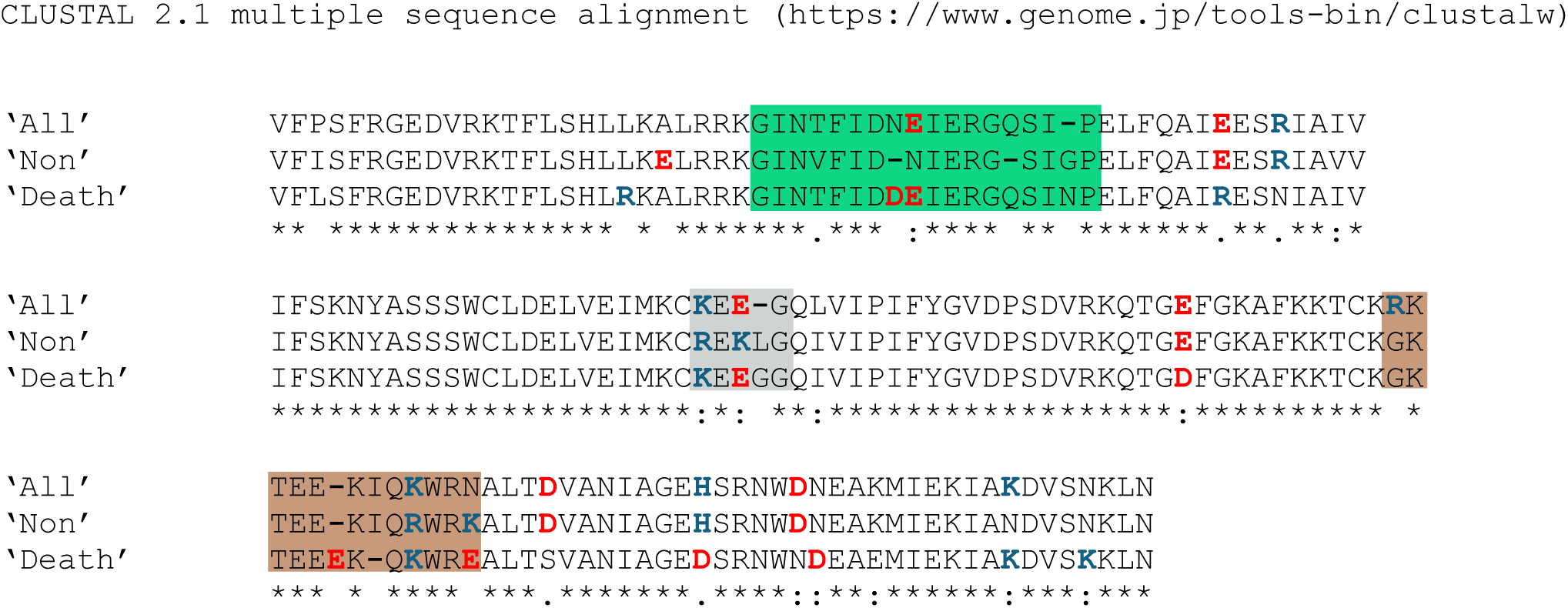
ClustalW alignment of consensus artificial TIRs. Alignment of the three artificial TIRs, ‘Death’, ‘All’, and ‘Non’. Alignment performed with ClustalW (https://www.genome.jp/tools-bin/clustalw). Asterisk indicates residue conservation. Positive and negative charged residue variations are shown blue and red, respectively.

**SI 9.**
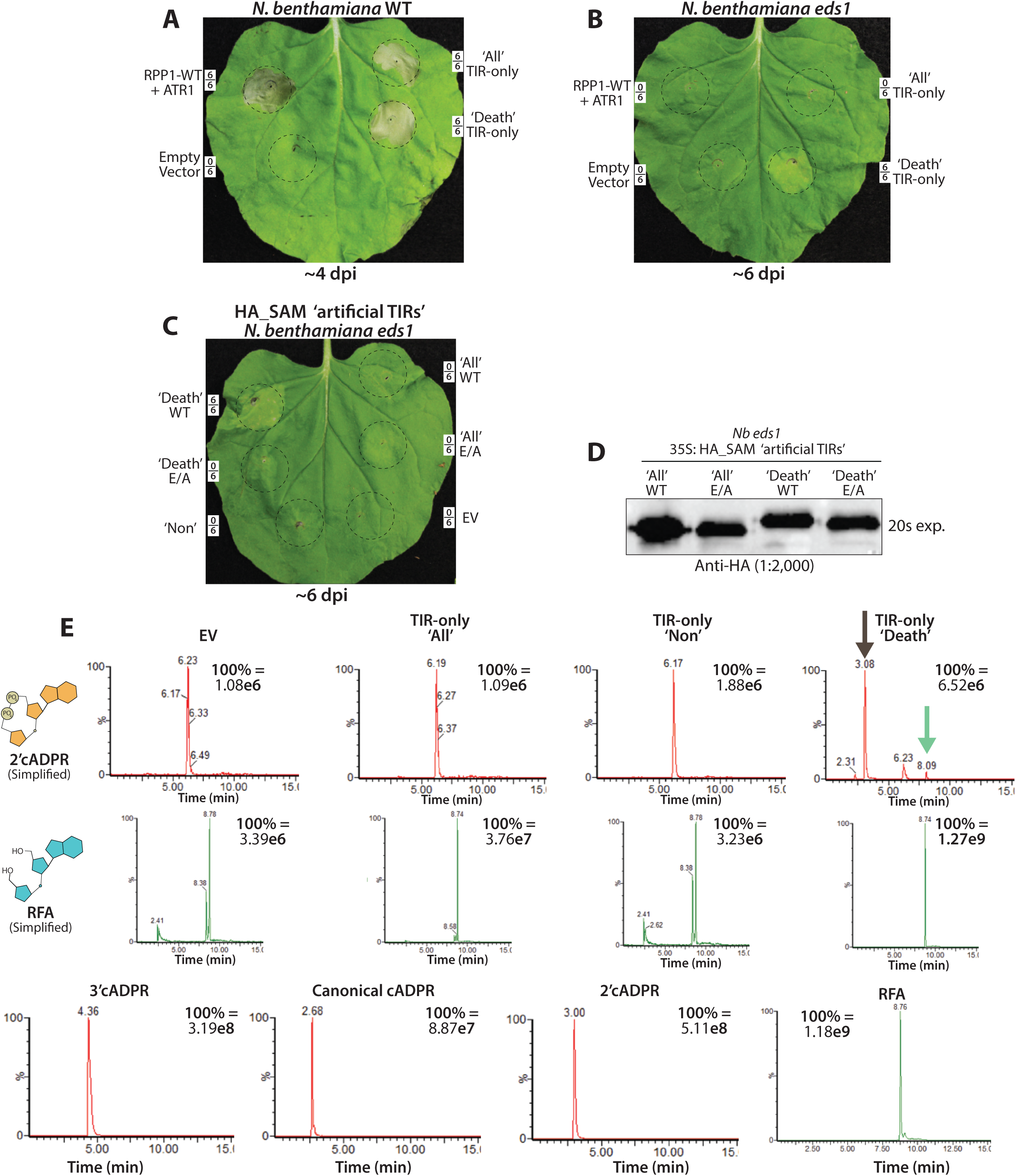
Artificial TIRs signal cell death and generate metabolites as TIR-only proteins; ‘Death’ causes chlorosis in Nb eds1. (*A*, *B*) *Nb* WT or *eds1* leaves expressing TIR-only versions of the artificial consensus TIRs, ‘Death’, ‘All’ or ‘Non’. Negative control empty vector (35S:YFP), positive control RPP1-TNL (WsB) co-expressed with ATR1 (Emoy) effector. (*C*) As in A and B, but showing *Nb eds1* leaves expressing various HA-SAM tagged artificial TIRs. (*D*) Anti-HA immunoblot of HA_SAM artificial TIR catalytic glutamate (E to A, alanine) mutants. (*E*) LC-MS chromatographs of products of artificial TIRs (TIR-only proteins expressed in *Nb eds1*), relative to standards. A black arrow denotes 2’cADPR peak, and a green arrow indicates an unknown 542 Da (Dalton) peak with ∼8.10 min retention time.

**SI 10.**
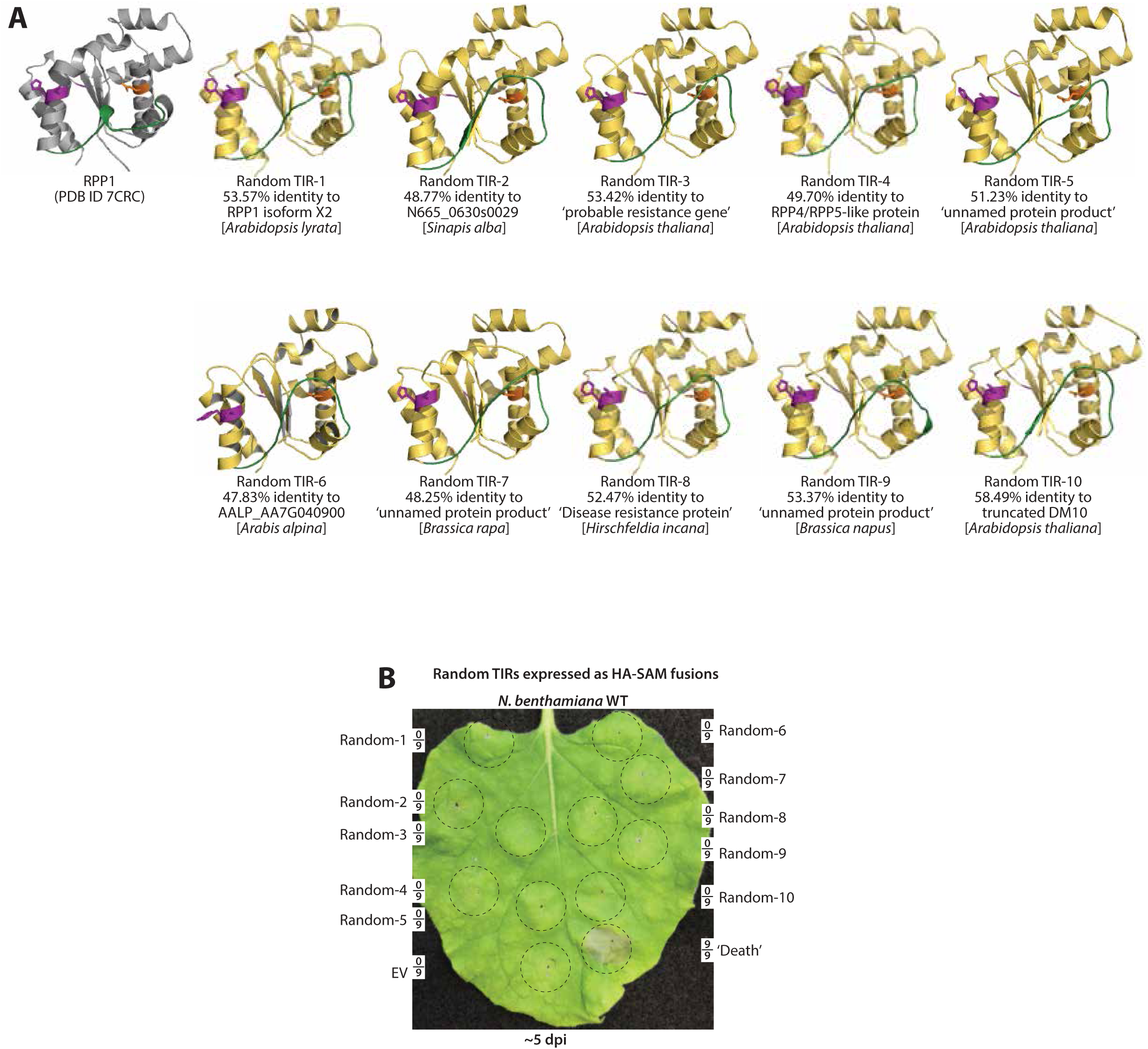
Examined ‘random’ artificial TIRs do not signal cell death in N. benthamiana. (*A*) RPP1 TIR (active state; PDB 7CRC) structure and AlphaFold2 predictions of randomly designed artificial TIRs. ‘Random TIRs’ were designed by randomly selecting amino acids (in linear order) from 146 Arabidopsis TIRs (Two xTNx TIRs were not included). The closest natural TIR (and percent identity) is listed below the structure models. (*B*) *Nb* WT expressing random artificial TIRs as HA_SAM fusions at OD 0.80. EV (35S:GFP, empty vector). Leaves imaged ∼4-5 dpi, and framed numbers denote replicates per set.

**SI 11.**
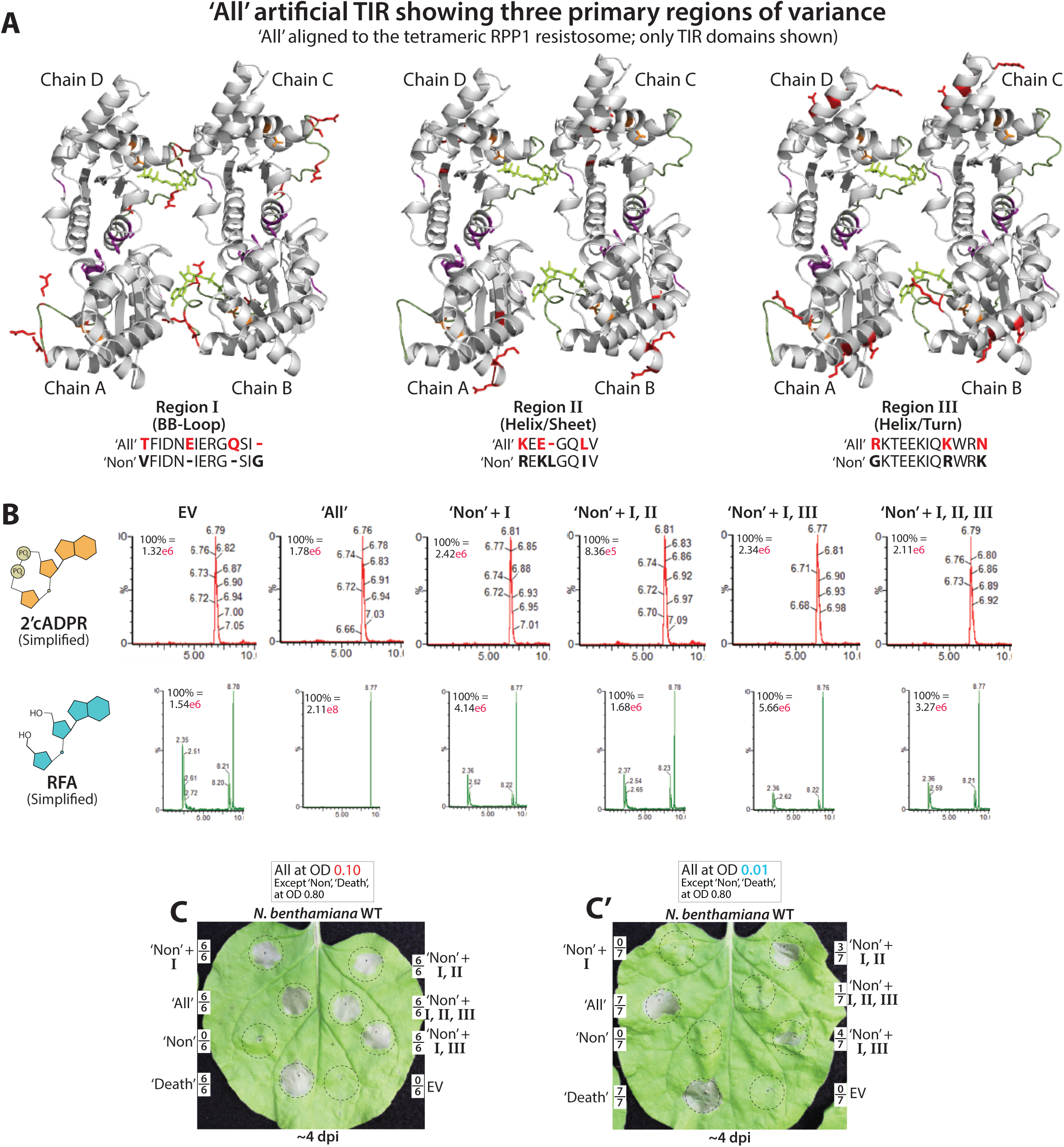
Variation at regions II and III does not increase the activity of ‘Non’ + the BB-loop of ‘All’. (*A*) The ‘All’ consensus TIR (AlphaFold2) modeled onto the tetrameric structure of the RPP1 TIR-NLR resistosome (PDB: 7CRC). Residues variant between ‘All’ and ‘Non’ are shown in red and defined as region I, II, or III, respectively. Region I corresponds to the BB-loop, and regions II and III positions to helices αC and αD, respectively. The TIR interfaces are shown purple, catalytic glutamate (E) in orange, and the BB-loop and ATP are shown in green. (*B*) LC-MS chromatographs of 2’cADPR and RFA from ‘Non’, replacement variants of ‘Non’, or ‘All’, expressed in *Nb eds1* leaves. (*C*, *C*’) *Nb* WT leaves expressing ‘Non’, replacement variants of ‘Non’, or ‘All’. The regions transferred from ‘All’ into ‘Non’ are noted above in Panel A. Constructs in Panel C were delivered at OD_600_ 0.10 while those in C’ were at OD_600_ 0.01. EV: empty vector (35S:YFP). Leaves were imaged ∼4-5 dpi, and framed numbers denote replicates per set.

**SI 12.**
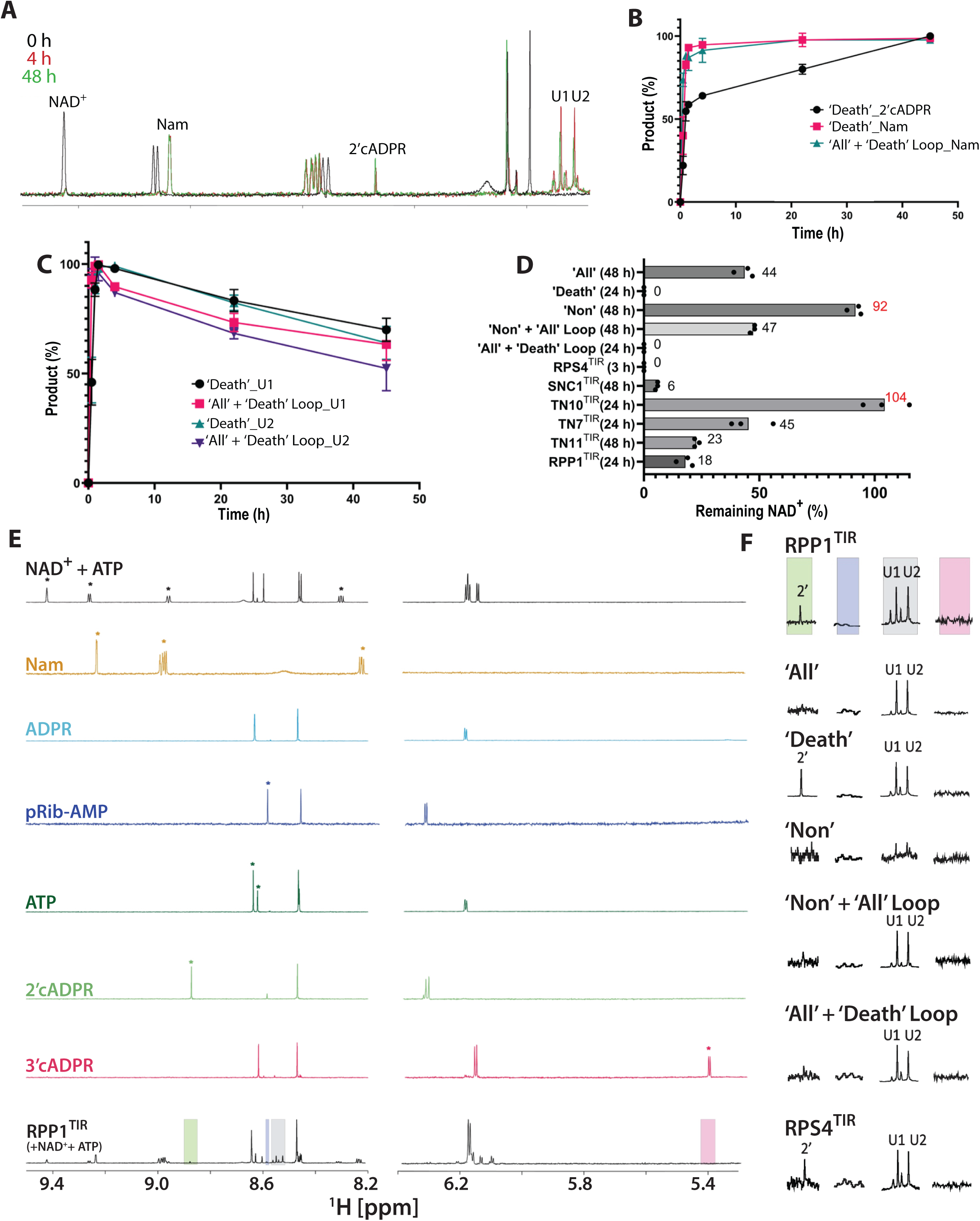
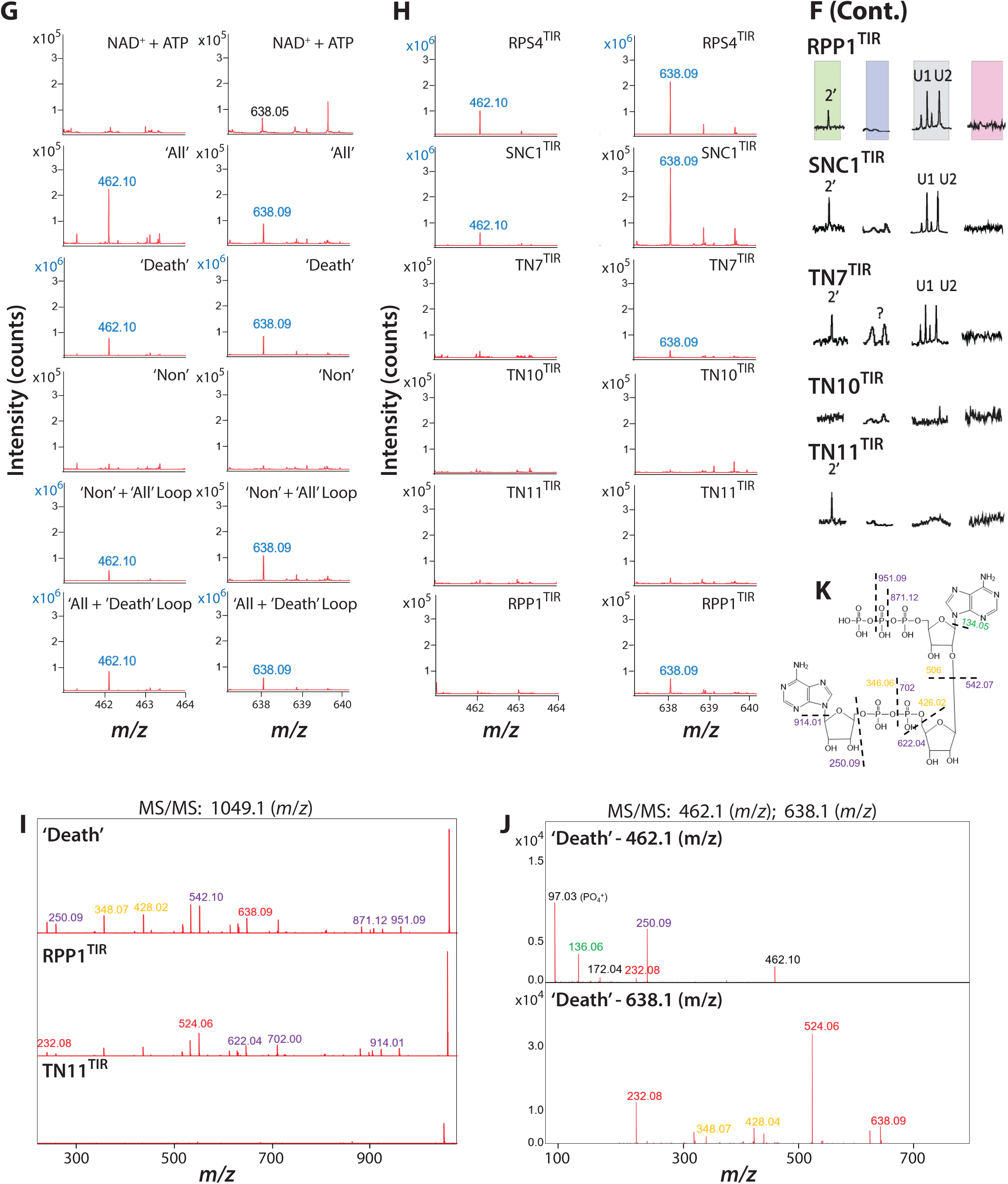
NMR and MS analysis of *in vitro* TIR NADase reactions. (*A*) Representative ^1^H NMR spectra from ‘Death’ TIR samples collected at 0 h, 4 h and 48 h. Peaks labelled with NAD^+^, Nam (nicotinamide), 2’cADPR, U1 and U2 were used for quantification. (*B*, *C*) NMR quantification of Nam, 2’cADPR, and U1 and U2 for ‘Death’ and the ‘All’ + ‘Death’ Loop TIRs. Percent intensity calculated relative to the time-point with the highest intensity. (*D*) Percent NAD^+^ remaining from endpoint NADase assays of natural and artificial TIR domains. Incubation time of each TIR indicated by brackets. Red text indicates no NAD^+^ cleavage and no Nam buildup; *i.e*. inactive TIRs. (*E*) ^1^H NMR spectrum of individual standards (∼50 μM) NAD^+^, ATP, Nam, ADPR, 2’cADPR, 3’cADPR and pRib-AMP compared to the spectrum of completed NADase reaction of RPP1^TIR^. Asterisks indicate peaks used to identify compounds. Colored boxes in RPP1 TIR spectra indicate regions aligning to particular metabolites: 2’cADPR (green), unique peaks (U1 and U2, grey), pRib-AMP (blue). (*F*) Region of ^1^H NMR spectrum used for NADase reaction monitoring of assayed artificial and natural TIRs. (*G*, *H*) MALDI-MS showing additional unknown species at 638.09 and 462.1 m/z. Spectra with blue text confidently detect these unknown species. (*I*, *K*) MS/MS secondary fragmentation of 1049.06 (ADPr-ATP) from ‘Death’, RPP1^TIR^ and TN11^TIR^ (no ADPr-ATP detected) samples. Purple text indicates peak fragments with a positive charge, while yellow indicates negative fragments (gain 2H to acquire a +1 charge). Red text fragments are unidentified but were also observed in 462.1 and 638.1 *m/z* MS/MS data. (*J*) MS/MS fragmentation of 462.1 and 638.1 m/z compounds from ‘Death’ TIR. MS tune settings were slightly adjusted when fragmenting the compound 461.1 m/z, to detect smaller fragments (see Methods).

**SI 13.**
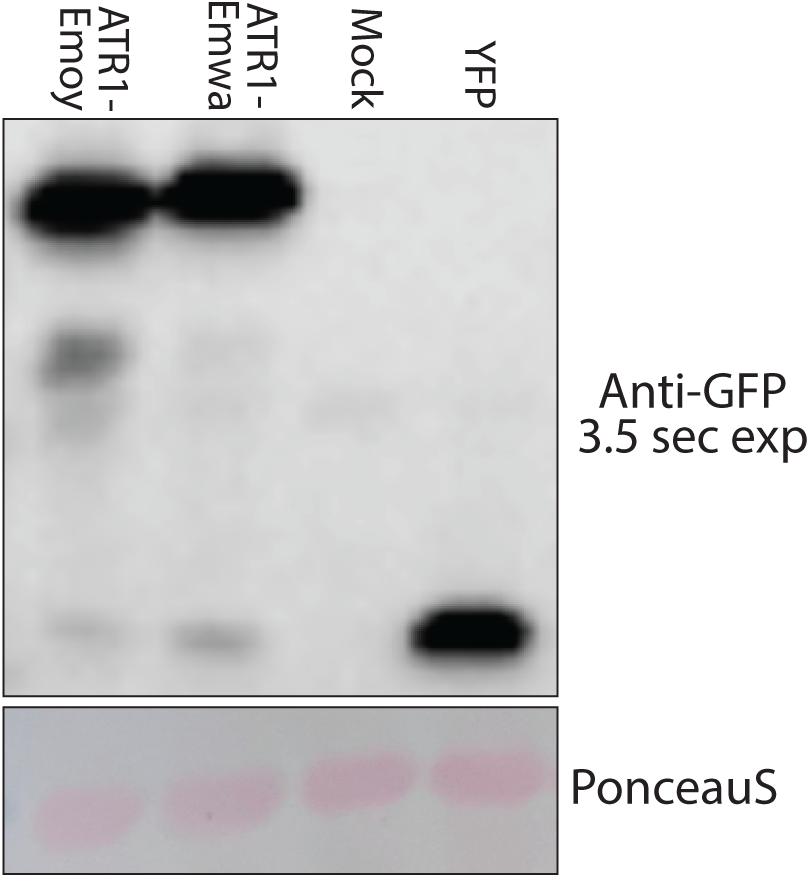
Accumulation of ATR1 proteins. Anti-GFP immunoblot of lysates from *N. benthamiana eds1* leaves expressing C-terminal YFP-tagged ATR1 Emoy or Emwa. Samples harvested 48 hpi agroinfiltration. ATR1-Emoy activates the RPP1 WsB allele utilized in this study. Positive control 35S:YFP, negative control mock (buffer) infiltrated leaves. PonceauS stain indicates relative protein abundance within each sample.

